# Sterile Soil as Prebiotics Mitigate Intergenerational Loss of Gut Microbial Diversity and Anxiety-Like Behavior Induced by Antibiotics in mice

**DOI:** 10.1101/2023.04.15.536315

**Authors:** Na Li, Xiaoao Xiao, Honglin Zhang, Zhimao Bai, Mengjie Li, Jia Sun, Yangyang Dong, Wenyong Zhu, Zhongjie Fei, Xiao Sun, Pengfeng Xiao, Yuanqing Gao, Dongrui Zhou

## Abstract

The decline in gut microbial diversity of modern human is closely associated with the rising prevalence of various diseases. It is imperative to investigate the underlying causes of gut microbial loss and the rescue measures. Although the impact of non-perinatal antibiotic use on gut microbiota has been recognized, its intergenerational effects remain unexplored. Our previous research has highlighted soil in the farm environment as a key prebiotic for gut microbiome health by restoring gut microbial diversity and balance. In this study, we investigated the intergenerational consequences of antibiotic exposure and the therapeutic potential of soil prebiotics. We treated mice with vancomycin and streptomycin for 2 weeks continuously, followed by 4-8 weeks of withdrawal period before breeding. The process was repeated across 3 generations. Half of the mice in each generation received an oral soil prebiotic intervention. We assessed gut microbial diversity, anxiety behavior, microglia reactivity, and gut barrier integrity across generations. The antibiotics exposure led to a decrease in gut microbial diversity over generations, along with aggravated anxiety behavior, microglia abnormalityies, and altered intestinal tight junction protein expression. Notably, the third generation of male mice exhibited impaired reproductive capacity. Oral sterile soil intervention restored gut microbial diversity in adult mice across generations, concomitantly rescuing abnormalities in behavior, microglia activity, and intestinal barrier integrity. In conclusion, this study simulated an important process of the progressive loss gut microbiota diversity in modern human and demonstrated the potential of sterile soil as a prebiotic to reverse this process. The study provides a theoretical and experimental basis for the research and therapeutic interventions targeting multiple modern chronic diseases related to intestinal microorganisms.

## Introduction

With the advent of industrialization, the diversity of human intestinal microbiota is progressively declining from one generation to another [1-5]. It is imperative to investigate the underlying causes contributing to this loss of gut microbial diversity? [4, 6, 7]. Antibiotics, indispensable in modern medicine, primarily have broad-spectrum activity and disrupt the gut microecological balance while eliminating pathogenic bacteria [6]. The diversity of gut microbiota in mice is linked to the microglia activity and neurodevelopment process [8]. Additionally, antibiotics have been found to induce damage to the intestinal barrier [9, 10], and affect social behavior, anxiety, and depression-like symptoms in mice [11-15].

Limited research exists on the transgenerational impact of antibiotic-induced gut microbial loss outside the perinatal period [6]. Erica et al. reported that a low microbiota-accessible carbohydrate diet across four successive generations can exacerbate the loss of gut microbial diversity [2]. Reintroduction of a microbiota-accessible carbohydrate diet failed to fully restore gut microbial diversity [2]. Blaser’s clinical observations suggest that gut microbes in modern society are disappearing generation by generation, with antibiotics being the primary culprit, warranting urgent measures to repair gut microecology [6].

Numerous studies have focused on the restoration of gut microecology through probiotics, prebiotics, and fecal microbiota transplantation [16-18]. Recent research has highlighted the influence of a low cleanliness living environment on gut microbial composition and diversity [19-23]. The diversity of gut microbiota in mice living in a low-cleanliness environment is higher than those in a high-cleanliness environment [20]. Furthermore, the gut microbial diversity of mice living in a low cleanliness environment rapidly recovers after antibiotic exposure, while the recovery of mice living in a high cleanliness environment is slower [20]. What factors affect gut microbiota structure and composition in a low cleanliness living environment? Zhou et al. found that soil plays a key role in shaping intestinal microbial structure in low cleanliness environments [24, 25]. Soil can act as a “prebiotic,” providing airborne microorganisms that, when consumed, increase intestinal microbial diversity and ecological balance in mice [24]. However, it remains to be determined whether soil prebiotics can effectively restore antibiotic-induced intestinal microecology imbalance.

In this study, we aimed to determine whether antibiotics induce a stepwise extinction of gut microbiota and whether this leads to exacerbation of anxiety behavior and intestinal functional barrier disruption in mice. Additionally, we explored the potential of soil intake as a prebiotic to restore the antibiotics-induced intestinal microecology imbalance, anxiety behavior, and intestinal functional barrier in mice by concurrently inhaling airborne microbes. In our experiment design and process, we used mouse living in SPF house to simulate the human lifestyle. Although the cleanliness level in the SPF animal room was higher than that of modern human living environments, it facilitated rapid modeling of changes in intestinal microecology and the pattern identification. We used metal mesh to separate mouse feces, simulating fecal-oral separation in humans. Open cages were allowed for modeling of microbial communication among humans. Soil rescue was performed on mice in a general animal room, also simulating human living environment. The air in humans living environments contains numerous microorganisms, which are constantly inoculated into our bodies through respiration or diet.

## Results

### 1. Antibiotic stepwise decreased gut microbial diversity and function genes abundance

Parental mice were exposed to antibiotics for three generations to obtain offspring model mice (Fig. 1A). High-throughput sequencing of microbial 16S rDNA from fecal samples revealed a generational reduction in the number of operational taxonomic units (OTUs) in mice gut microbiota of mice (Fig. 1B). This trend was further confirmed by the Chao1 index (Fig. S1 and Table S1). Principal coordinate analysis (PCoA) of unweighted unifrac distances indicated that the gut microbiota of G1, G2, and G3 mice progressively diverged from the control G0 mice (Fig. 1C and 1D). Cluster analysis of the top 200 OTUs based on their total abundance showed that some OTUs of the gut microbiota were reduced in G1 mice compared to G0 mice, and this number increased in G2 and G3 groups (Fig. 1E and Table S2).

**Figure 1.**
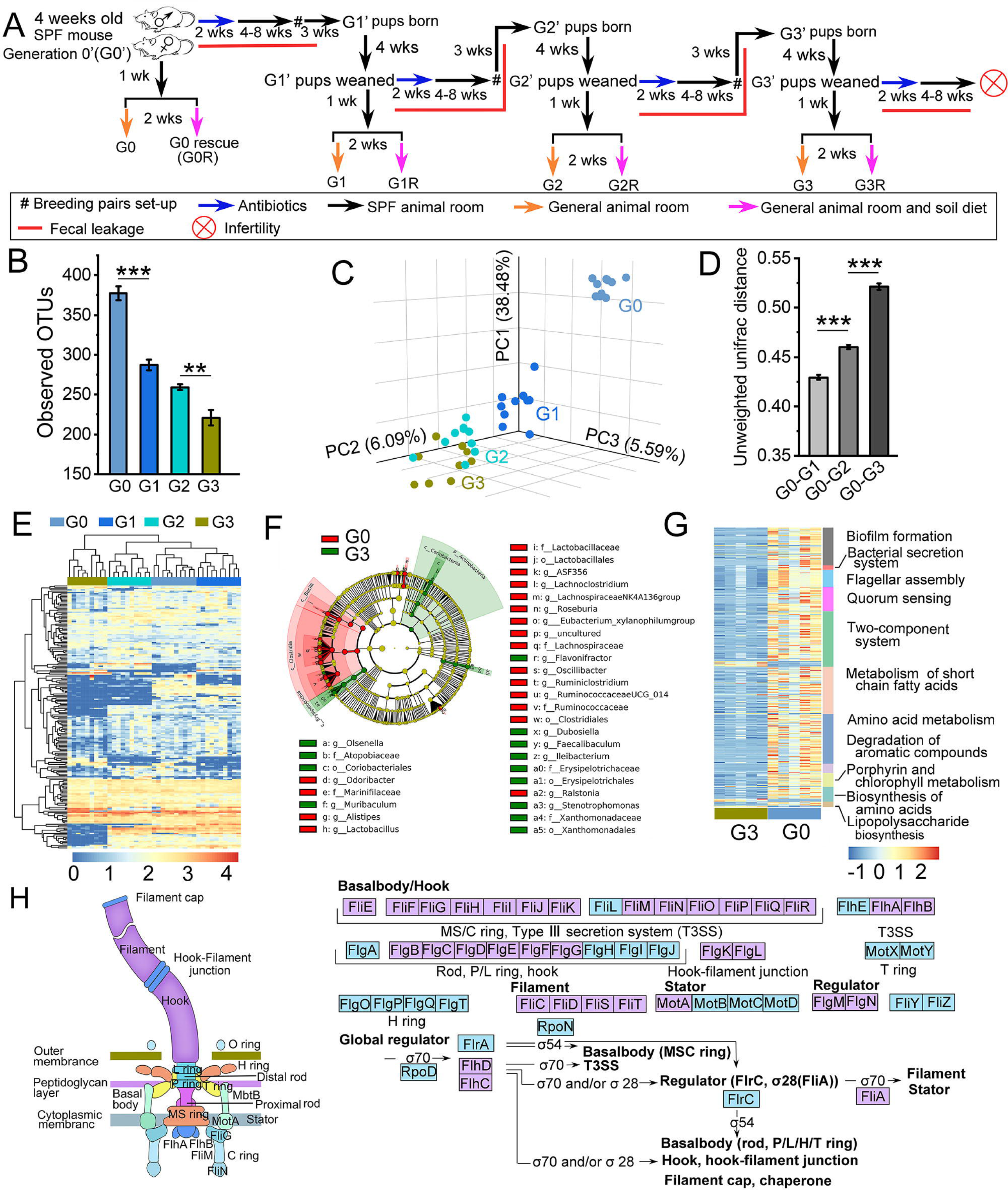
Effects of Antibiotic Treatment on Gut Microbiota in Three Generations of Mice. A. Schematic of the mouse model used in the study. SPF mice (G0) were treated with antibiotics for 2 weeks and were allowed to reproduce 4-8 weeks after stopping the medication. The offspring were designated as G1, and the process was repeated to obtain G2 and G3. Male G3 mice lost their ability to reproduce further. The breeding process was conducted in the SPF animal room, and feces were leaked for adult mice. Mice were taken to a general animal room to soil intake intervention as prebiotics. B. Number of unique operational taxonomic units (OTUs) at a 97% threshold of fecal microbiota. C. Principle coordinates analysis (PCoA) for unweighted unifrac distance of 16S rRNA gene sequences. D. Comparison of unweighted unifrac distance of 16S rRNA gene sequences between each generation and G0 mice. E. Heat map profile of top 200 OTUs with total abundance. Color represents OTU abundance [lg (abundance of OTUs + 1)]. F. Cladogram generated by LEfSe indicating differences in taxa between G0 and G3 groups. Each successive circle represents a phylogenetic level (phyla, class, order, family, genera). G. Heat maps of functional genes compared between G3 and G0 groups. The color shows the Z score of relative abundance. H. Functional genes associated with flagellar assembly. The purple box represents G0 highly expressed genes, and the blue box represents genes without a difference or undetected. B,D: Error bars represent the standard error of the mean (SEM), and P values are based on ANOVA with Bonferroni post-hoc test. *P < 0.05, **P < 0.01, ***P < 0.001. C-F: N=9-10. G-H: The p-value of the Mann-Whitney U test is greater than or equal to 0.05, or the fold change of two groups is less than 2. N=5.

To further explore the differences in the gut microbiota between the G3 and G0 groups, we performed linear discriminant analysis effect size (LEfSe) analysis of 16S rRNA gene sequences (Fig. 1F, S2 and S3). This analysis identified a significant decrease in thirteen genera, including Odoridacter and Lachnoclostridium, and a significant increase in seven genera, such as Ileibacterium and Olsenella, in the G3 group compared to the G0 group (Fig. 1F, S2 and S3).

To further investigate the impact of microbial loss on gut microecological functions in mice, we conducted shotgun sequencing analysis of gut microbes in G0 and G3 groups. Our results revealed a significant reduction in the number of functional genes in the G3 group compared with the G0 group. Specifically, major functional genes related to biofilm formation, bacterial secretion system, flagellar assembly, quorum sensing, two-component system, metabolism of short chain fatty acids, amino acid metabolism, degradation of aromatic compounds, porphyrin and chlorophyll metabolism, biosynthesis of amino acids, and lipopolysaccharide biosynthesis were diminished in the G3 group (Fig. 1G and Table S3). Notably, among the 42 flagellar assembly-related genes detected, 33 genes were significantly different between the two groups, and the abundance of all 33 genes was lower in G3 mice compared to G0 mice (Fig. 1H and Table S3). Moreover, some functional genes associated with metabolism of butyric acid, the bacterial secretion system, and two-component system were significantly reduced in the G3 group (Fig. S4, S5, S6 and Table S3). Overall, these findings demonstrate that the loss of many gut microbes was associated with a decline in functional genes.

### 2. Soil rescue increased gut microbial diversity and function genes abundance in adult mice

The study analyzed the effect of soil rescue on gut microbial diversity. Soil rescue was found to significantly increase (P<0.05) the gut microbial Chao 1 estimate in the G1R, G2R, and G3R groups compared to the G1, G2, and G3 groups, respectively. However, there was no significant difference between the G0R and G0 groups (Fig. 2A). PCoA of unweighted unifrac distances revealed that soil rescue shifted the gut microbial composition of each generation of mice towards the G0 group (Fig. 2B-F). Moreover, soil rescue reduced the distance between the gut microbiota of each generation of mice and the G0 group (Fig. 2G). Further analysis of the G0, G0R, G3, and G3R shotgun sequencing using PCoA of Bray-Curtis distance also demonstrated a trend of gut microbiota in the G3R group moving towards the G0 group (Fig. S7C).

**Figure 2.**
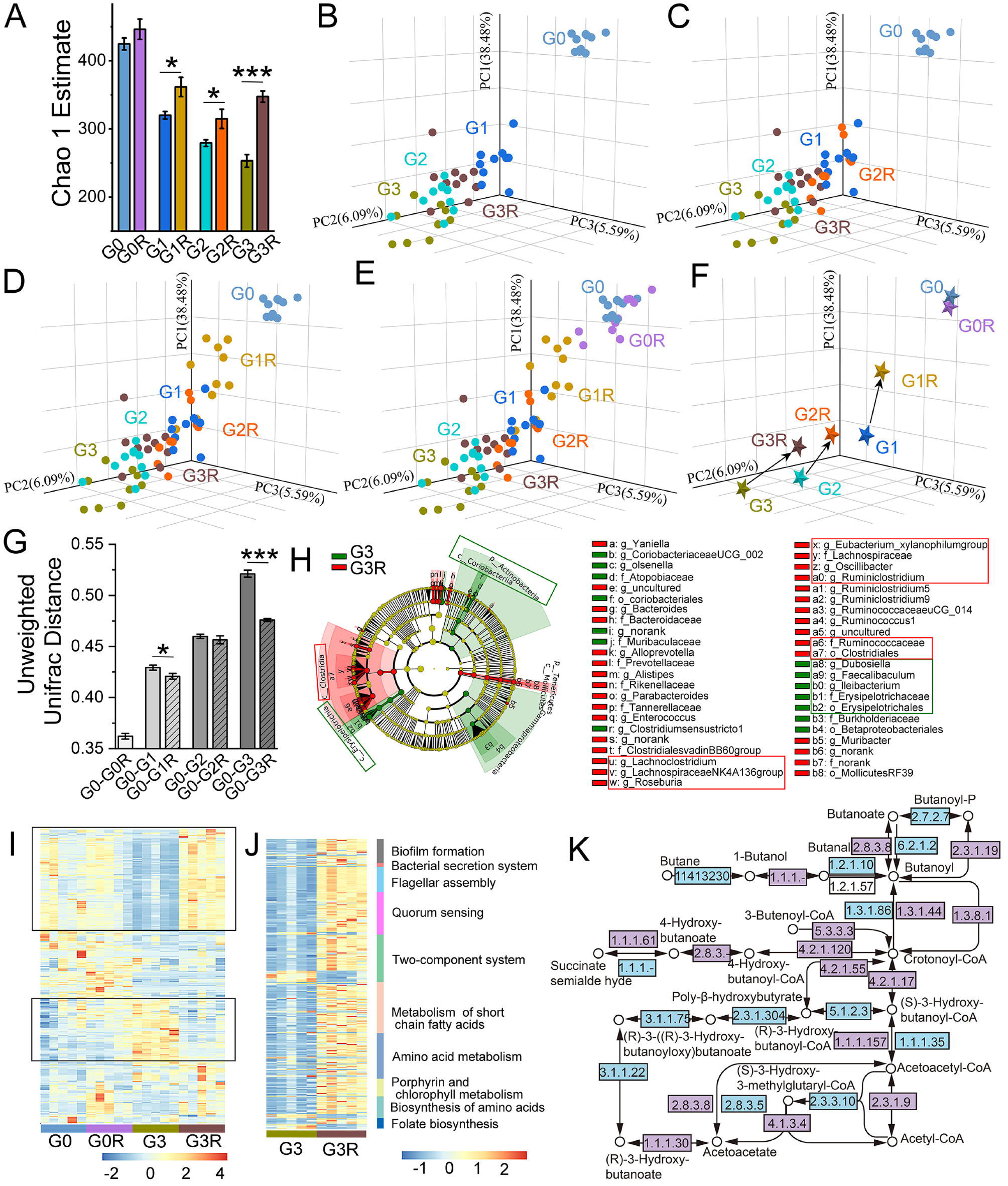
Comparative Analysis of Soil Rescue on Gut Microbial Function Restoration. A. Chao 1 estimate. B-E. PCoA plots of unweighted unifrac distance of 16S rRNA gene sequences. F. Change in PCoA trend after soil rescue for each generation. G. Comparison of unweighted unifrac distance of 16S rRNA gene sequences. Error bars represent s.e.m, and P-values were calculated based on a 2-tailed t-test (*P < 0.05, ***P < 0.001). H. Cladogram generated by LEfSe indicating differences in taxa between group G3 and group G3R. The red boxes outline the taxa that were significantly reduced in abundance in the G3 group compared to the G0 group and significantly recovered after soil rescue in the G3R group mice. The green boxes outline the taxa that were significantly increased in abundance in the G3 group compared to the G0 group and significantly recovered in the G3R mice. I. Heat map profile of the top 500 species with total abundance. J. Heat maps of functional genes compared between G3 and G3R groups. The color represents the Z-score of relative abundance. K. Functional genes associated with butyric acid metabolism. The purple box represents highly expressed genes in G3R, and the blue box indicates genes with no significant difference or were undetected. A-H. N=9-10; I-K. N=5. The P-value of the Mann-Whitney U test is greater than or equal to 0.05 or the fold change of two groups is less than 2.

Based on the analysis of alpha and beta diversity, it is evident that G1, G1R, G2, and G2R are in the intermediate transition stage. For further investigation of the changes in gut microbial function after three generations of antibiotic treatment and the effect of soil rescue, we selected G0/G0R and G3/G3R for analysis. These two generations and their intervention mice were also chosen for research in the later study of newborn mice.

LEfSe results of the previous 16S rRNA gene sequences revealed a significantly lower abundance of multiple bacteria in the G3 group of mice compared to the G0 group (Fig 1F and S2). However, the abundance of most bacteria among them was significantly higher in the G3R group than the G3 group (Fig. 2H, S8 and S9). Some taxa decreased in the G3 group and increased in the G3R group following soil rescue, as shown in the red boxes in Fig. 2H. Additionally, there were taxa that increased in the G3 group and recovered after soil rescue, as shown in the green boxes in Fig. 2H (Fig. 1F, S2, S3, 2H, S7, and S8). Species abundance analysis of the shotgun sequences annotation further verified these results (Fig. 2I and Table S4).

Similarly, functional genes lost in the G3 group were partially restored in G3R mice after soil rescue. These genes included those related to biofilm formation, bacterial secretion system, flagellar assembly, quorum sensing, two-component system, metabolism of short-chain fatty acids, amino acid metabolism, porphyrin and chlorophyll metabolism, biosynthesis of amino acids, and lipopolysaccharide biosynthesis (Fig. 2J, 2K, S10, S11, S12 and Table S5). Among the 41 flagellar assembly-related genes detected, 32 genes showed significant differences between the two groups, with greater abundance in G3R than in G3 (Fig. S10 and Table S5).

In summary, soil rescue intervention significantly increased the diversity of gut microbiota in antibiotic-treated mice offspring and shifted the microbial composition of each generation towards the G0 mice profile. Furthermore, soil rescue resulted in a substantial recovery of functional genes, particularly those related to flagellar assembly and butyric acid metabolism.

### 3. Soil rescue alleviated anxiety-like behavior and microglial abnormality in adult mice

We conducted further analysis on anxiety-like behavior and explored the potential mechanisms involving gut-brain axis in G0/G0R and G3/G3R mice. The open-field test results showed that the duration and frequency of mice in the G3 group entering the open field center were significantly lower than those in the G0 group (Fig. 3A). In contrast, the duration and frequency of mice in the G3R group entering the open field center were significantly higher than those in the G3 group (Fig. 3A). Similar results were observed in the elevated plus-maze test (Fig. S13).

**Figure 3.**
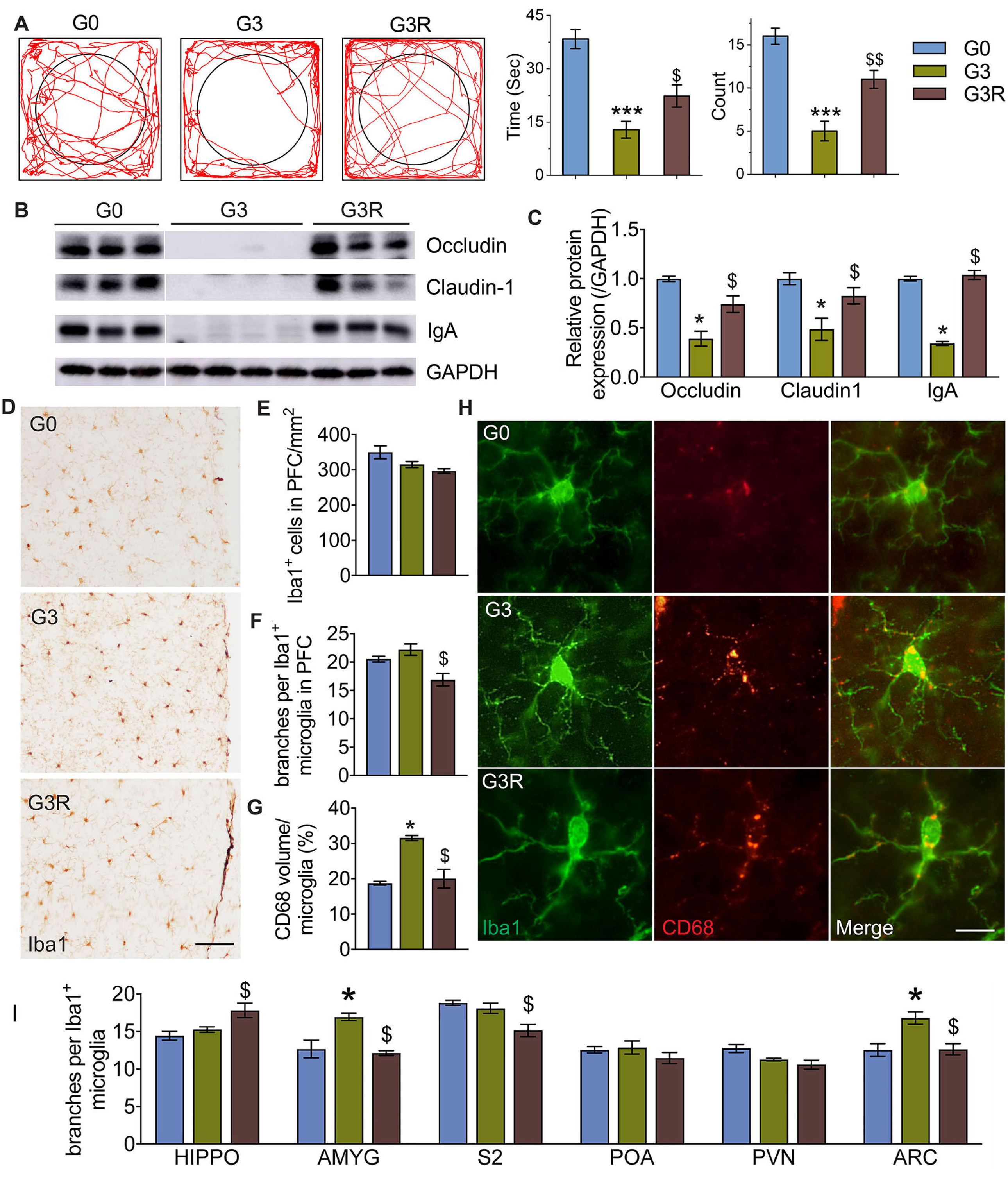
Effect of antibiotics and soil rescue on anxiety behavior, intestinal barrier, and microglia in the brain of G3 mice. A. Open-field test. B. Western blot showing the levels of Occuludin, Claudin-1, and IgA in the colon. C. Quantification of B. D. DAB staining with anti-Iba1 antibody in the prefrontal cortex region. E. Quantification of Iba1-ir positive cell number in D. F. Quantification of branches of individual microglia in D. G. Quantification of CD68-ir positive volume in individual microglia in D. H. Colocalization of CD68 and Iba1 in the prefrontal cortex region. I. Quantification of branches of individual microglia in each brain region. N=4-5 animals per group. Scale bar=100μm in D, Scale bar=25μm in H. *P<0.05 vs G0; $P<0.05 vs G3. All mice in this figure are male.

The western blot results showed that the levels of occludin, claudin-1, and IgA in the colon of the G3 group were significantly lower than those in the G0 group. Soil rescue significantly increased the levels of the three proteins in the intestine of G3R group (Fig. 3B and 3C). Microglia activity is crucial for synapse plasticity in the brain and influenced by gut microbiota, making them as a potential critical link between gut microbiota and animal behaviors. Therefore, we assessed the number and morphology of microglia in the brain using an anti-Iba1 antibody and evaluate the phagocytosis activity of microglia by colocalization of CD68 and Iba1. The results showed that there was no significant change in the number of microglia in G3 group (Fig. 3D, 3E and S14A). The G3 group had a significant increase in the number of microglia branches in the amygdala (AMYG), arcuate nucleus (ARC) brain regions compared to the G0 group (Fig. 3D, 3F, 3H, and 3I). Compared with G3, the microglia branches of G3R in the prefrontal cortex (PFC), amygdala (AMYG), secondary somatosensory cortex (S2), and ARC brain regions were significantly reduced (Fig. 3D, 3F, 3H, and 3I). The CD68-ir positive volume in individual microglia of PFC, AMYG, and ARC brain regions in G3 group was significantly higher than that in G0 group (Fig. 3G, 3H, and S14B). Compared with G3, the microglia activation in the FPC and AMYG brain regions of G3R decreased significantly (Fig. 3G, 3H, and S14).

In conclusion, these results indicate that soil rescue for adult mice significantly improved the intestinal barrier function, reduced microglia malformations, and attenuated the phagocytosis activity of microglia in anxiety-related brain regions.

### 4. Soil rescue of G3 infant mice restored gut microbiota nearly to the level of soil rescue of G3 infant mice

Based on the findings from the aforementioned studies, it was observed that soil rescue could partially restore some of the functions of gut microbiota in adult mice. Previous research has suggested that gut microbe colonization during early stages of life plays a crucial role in the development of the nervous and immune systems. [26-29]. To further examine the impact of soil rescue on the structure and function of gut microbiota in neonatal mice, we investigated the effect of soil rescue on infant mice on the second postnatal day (Fig. 4A). After 10 days of administering soil rescue, we sequenced the 16S rRNA genes of rectal microorganisms in infant mice, and randomly selected 5 infant mice from each group for metagenome sequencing.

**Figure 4.**
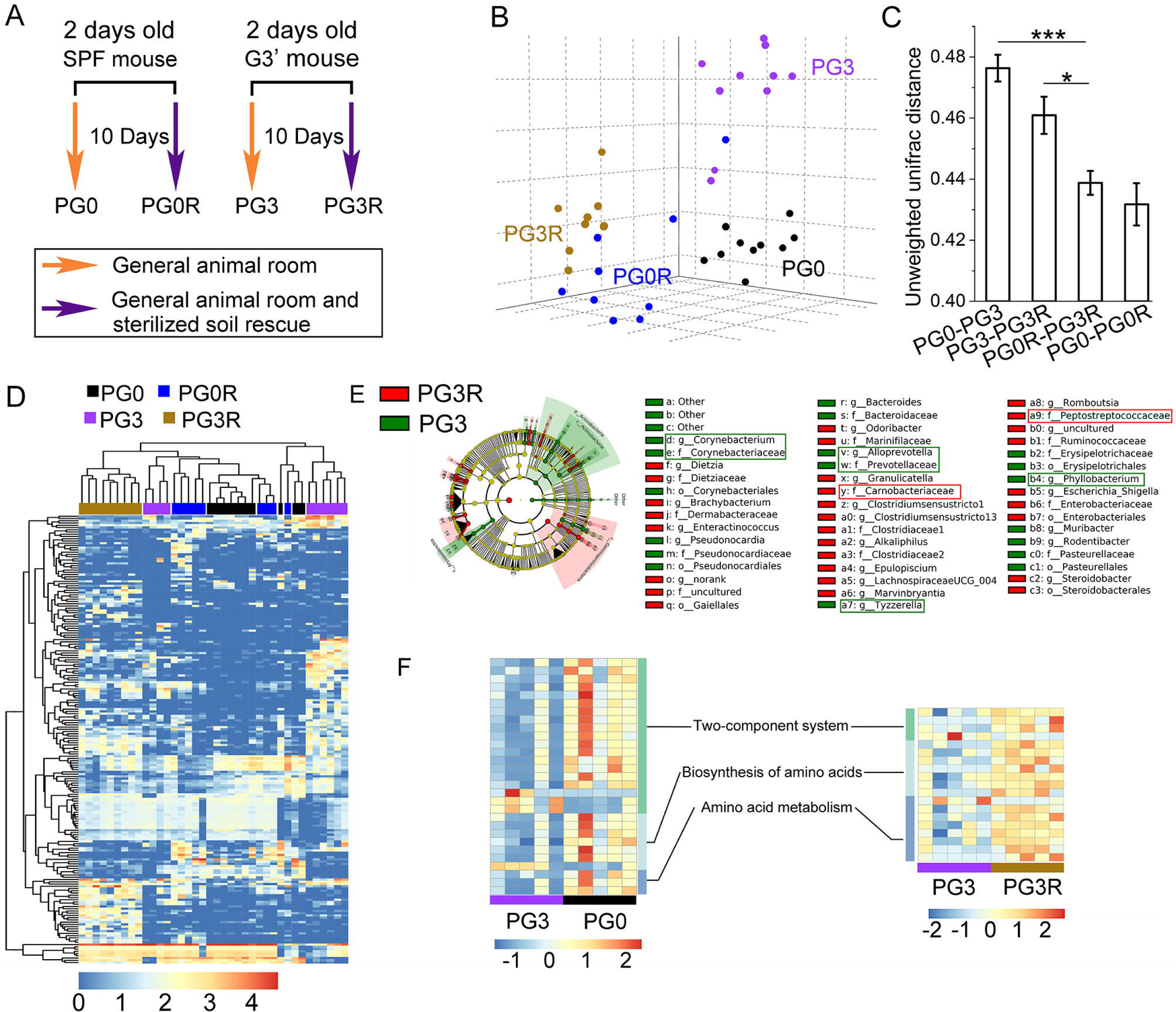
Effect of antibiotics and soil rescue on the gut microbiota of G3 infant mice. A. PG3R mice were fed with sterile soil from the second day of delivery to the 12th day, and some PG3R/PG0R mice were euthanized on the 12th day of age to collect tissue samples and rectal contents. Other PG3R/PG0R mice were raised in the general animal room until the age of 6 weeks for behavior testing. B. PCoA plot of unweighted unifrac distance of 16S rRNA gene sequences. C. Comparison of unweighted unifrac distance. Error bars represent s.e.m., and P values are based on ANOVA with Bonferroni post-hoc test. *P < 0.05, ***P < 0.001. D. Heat map profile of top 200 OTUs with total abundance. The color represents the abundance of OTUs [log(abundance of OTUs + 1)]. E. Cladogram generated by LEfSe. The red boxes outline the taxa that were reduced in abundance in the PG3 mice compared to PG0 mice and recovered after soil rescue. The green boxes outline those that increased in PG3 mice compared to PG0 mice and recovered after soil rescue. F. Functional genes with differences between PG3 and PG0 (left) and between PG3 and PG3R (right). The color shows the Z score of relative abundance, and P values are from the Mann-Whitney U test, where values less than 0.05 are considered significant.

The results indicated that the use of antibiotics in three consecutive generations of parent mice led to a significant difference in β diversity between PG3 and PG0 (Fig. 4B), with the longest unweighted unifrac distance between them (Fig. 4C). After soil rescue, the PG0R group was also separated from the PG0 group (Fig. 4B), but they remained the two nearest groups (Fig. 4B and 4C). From the column chart’s height, the next nearest groups were PG0R and PG3R (Fig. 4B). Thus, the gut microbiota composition of the PG3R group, was significantly closer to that of the PG0R mice. There was no significant difference between the PG0-PG0R and PG0R-PG3R distances, but the distance between PG0R-PG3R was significantly lower than that between PG3-PG3R (P<0.05, Fig. 4C), indicating a significant difference from the rescue effect of adult mice (Fig. 2G). Similar results were obtained on the heatmap of top 200 OTUs, namely, the gut microbiota of PG3R were closer to PGOR and PG0 (Fig. 4D and Table S6). In summary, soil rescue caused significant changes in PG3R mice compared to PG3 mice, bringing them closer to the gut microbiota composition of PG0 and PG0R mice.

We further analyzed the specific changes in gut microbiota of infant mice using LEfSe, and the results were similar to those of adult mice. This included taxa that significantly changed in the PG3 group compared to the PG0 group (Fig. S15 and S16) and recovered significantly after soil rescue (Fig. 4E, S17 and S18), as shown in the red and green boxes in Fig. 4E (Fig. 4E and S15).

The metagenome results showed that the species significantly different between PG0 and PG3 groups in the top 500 were all significantly higher in PG0 than in PG3 (Fig. S19A). After soil rescue, among the top 500 species, 12 species increased and three species decreased (Fig. S19B). Analysis of functional genes revealed that genes related to amino acid metabolism, amino acid biosynthesis, and two-component systems were significantly reduced in PG3 group compared to PG0 group, and the related functional genes were partially restored after soil rescue (Fig. 4F, Table S7 and Table S8)

In summary, after soil rescue from the second day after delivery, the intestinal microbial composition of PG3 mice tended to become similar to that of PG0 mice.

### 5. Soil rescue of G3 infant mice restored anxiety-like behavior and abnormal microglia nearly to the level of SPF mice

The results of the open-field test (Fig. 5A) and the elevated plus-maze test (Fig. S20) on the PG3R mice were similar to those of the G3R mice as mentioned in the previous results. Likewise, the intestinal barrier function was also comparable to the results of the adult rescue group (Fig. 5B, 5C). Interestingly, when we evaluated the microglia reactivity in the brain regions related to anxiety behavior in infant mice, the effect of soil rescue was surprising. The infant intervention mice showed significantly better outcomes than that of adult intervention mice. –

**Figure 5.**
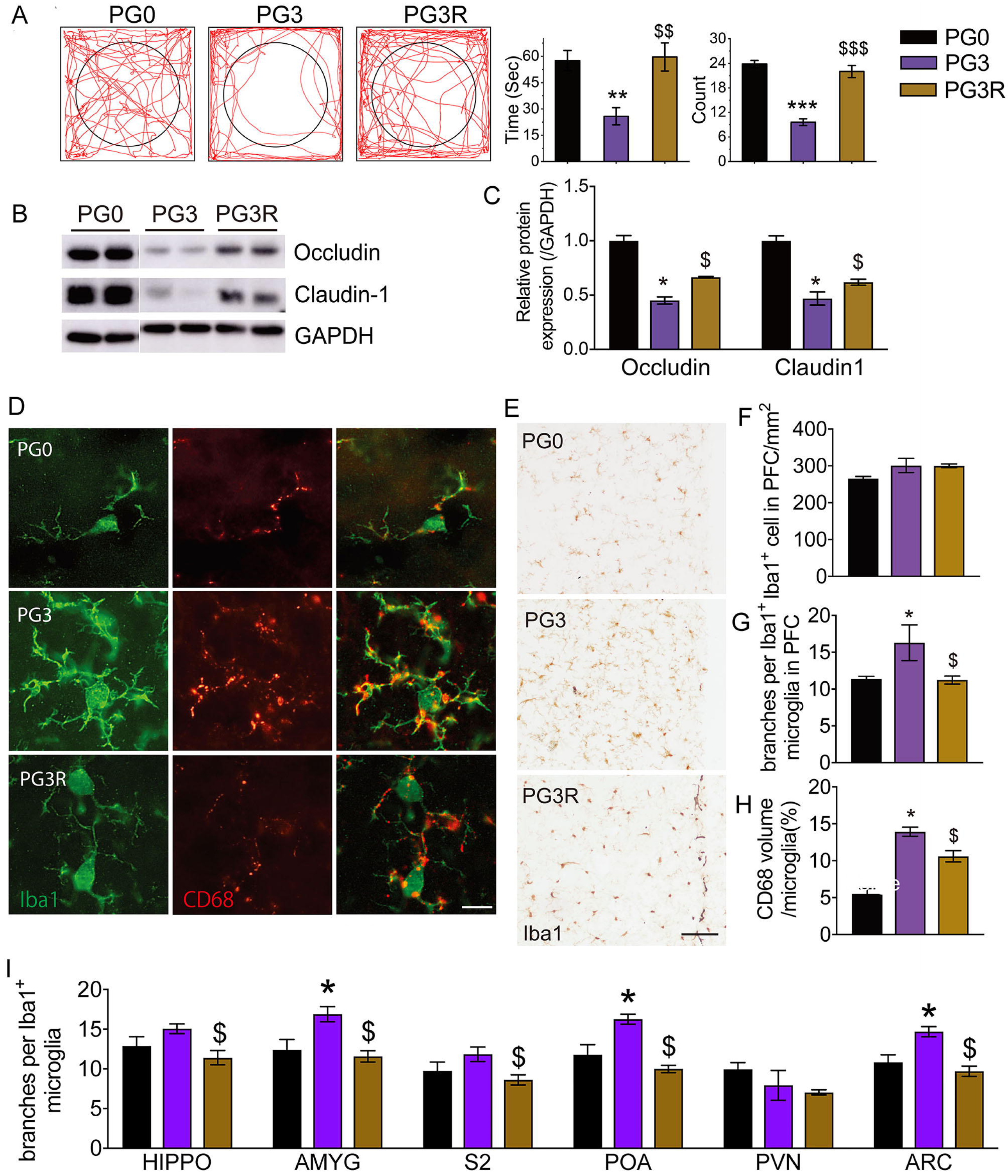
Effect of soil rescue on behavior, intestinal barrier, and microgilia for infant mice. A. Open-field test. B. Western blot of Occuludin, Claudin-1 and IgA of colon. C. Quantification of B. D. Colocalization of CD68 and Iba1 in prefrontal cortex region (PFC). E. DAB staining with anti - Iba1 antibody in PFC. F. Quantification of Iba1-ir postivie cell number in E. G. Quantification of branches of individual microglia in E. H. Quantification of CD68-ir positive volume in individual microglia in E. I. Quantification of branches of individual microglia in each brain region. N=4-5. Scale bar=100um in B, scale bar=25um in F. *P<0.05 vs PG0; $p<0.05 vs PG3. All mice are males.

PG3 infant mice exhibited evident microgliosis (with more branches) in most brain regions examined, including PFC, AMYG, preoptic area (POA), and ARC compared to the PG0 group (Fig. 5D, 5E, 5G, and 5I). The number of microglia in the hippocampus (HIPPO) was significantly lower in the PG3R group compared to the PG3 group (Fig. 4F and S21A). After soil intervention, the microgliosis were significantly attenuated in most brain regions, except for the paraventricular nucleus (PVN), and the morphology were comparable with PG0 in most brain regions (Fig. 5D, 5E, 5G, and 5I). The phagocytic activity of microglia (indicated by CD68 volume) showed a similar pattern that elevated in PG3 infant mice and alleviated by soil intervention in PG3R infant mice in most brain regions, especially in the PVN (Fig. 5D, 5H, and S21B).

Overall, initiating soil rescue from the second day after delivery resulted in the third-generation pups being more consistent with the control mice in terms of gut microbial structure, behavior, and brain microglia.

## Discussion

In this study, we investigated the effects of parental antibiotic use on offspring using SPF mice as an animal model. Our findings suggest that the gut microbiota diversity tended to decrease with each generation, and the third generation exhibited severe anxiety-like behavior, intestinal leakage, microglia malformation, and abnormal phagocytic activity. Soil rescue led to the restoration of various microorganisms and functional genes affected by antibiotic use, and the improvement gut microbial diversity intestinal leakage and the anxiely-like behaviors. Interestingly, soil rescue initiated from the 2nd day after delivery showed better repair function than intervention in adulthood, restoring all the indexes nearly to the level of SPF mice.

Our experimental results showed that sterile soil, like other prebiotics, promoted the growth of a large number of bacteria, and some bacteria that were inoculated from the air were able to survive with the support of the soil. However, environmental microorganisms could not colonize and survive in the intestinal tract of the control group without soil rescue, even if they lived in an environment containing a large number of microorganisms. Therefore, we chose the conventional animal room for soil rescue, which increased the differences between the soil rescue group and the control group, and better simulated the use environment of prebiotics in humans.

Antibiotic use reduces the gut microbial diversity of offspring mice generation by generation, even when antibiotics are used outside of the perinatal period. The experimental protocol used in this study simulated the routine clinical use of antibiotics in humans. In our experimental settings, all the mice were housed in a SPF animal room, and G1, G2, and G3 mice were in constant microbial exchange with G0 (SPF) mice through open cages. Despite this, the gut microbiota of mice still disappeared generation by generation after antibiotic exposure. It can be speculated that the disappearance of intestinal microorganisms caused by antibiotic treatment at any time in humans living in modern society will be transmitted to the next generation through vertical transmission. The increasing number of infants and children clinically suffering from allergic and autoimmune diseases [6, 30] may be related to the frequent use of antibiotics by their mothers. This study experimentally supports the hypothesis proposed by Martin, based on his clinical observations, that antibiotic use causes the gradual disappearance of human intestinal microorganisms [6].

Soil has a significant intervention effect on the gut microbiota in mice, which can replace the intervention of a low-cleanliness environment on the gut microbiota in animals. The gradual disappearance of human intestinal microorganisms is closely related to many chronic diseases, such as allergies, autoimmune diseases, obesity, and autism [31]. Therefore, finding effective techniques for intestinal microbial recovery is of great significance. Our previous study demonstrated that a low-cleanliness environment can rapidly restore antibiotics-damaged gut microbial diversity in mice [23]. However, low-cleanliness interventions have poor operability in clinical practice. This study demonstrates that soil, a key influence on gut microbiota in low-cleanliness environments [24], has a significant restorative effect on gut microbes that disappear generation by generation in mice.

The effectiveness of soil rescue was observed to be more pronounced in infant mice as compared to adult micein terms of gut microbial composition, microglial reactivity and animal behaviors. This could be attributed to the fact that early colonization of intestinal microorganisms is critical for gut microbial maturation and function [32]. Also, the first two weeks of postnatal life is a critical time window for synaptic pruning and neural circuit formation. The changes of microglial function during this period is strongly suggested in many maternal immune activations induced neurodevelopment diseases like autism [33, 34]. The changes in functional genes were not as prominent in infant mice as compared to adult mice, possibly due to the fact that the gut microecology of 12-day-old mice is still immature and undergoing substantial changes.

Soil rescue has been found to restore intestinal barrier integrity, microglial function, and behavior deficits in mice, which may be attributed to the restoration of short-chain fatty acid synthesis function. Our results showed that functional genes related to short-chain fatty acid metabolism were decreased with the loss of gut microbes in G3 mice. After intervention of soil, the abundance of short-chain fatty acid genes was restored in G3 mice, and the anxiety-like behavior, microglial function, and intestinal barrier function were significantly alleviated. Short-chain fatty acids have been reported to play a crucial role in regulating the central nervous system, with several studies highlighting their importance in microglial maturation [8, 35, 36]. Butyric acid, in particular, has been found to have antidepressant effects in animal studies and was shown to elevate concentrations of neurotrophic factor in the hippocampus and frontal lobes of the brain [37-40]. Furthermore, short-chain fatty acids have been found to regulate the intestinal functional barrier in several studies [41, 42].

Further experiments are needed to determine whether the restoration of bacterial secretion systems and flagellar assembly are essential for pathogenic bacteria to infect humans, and plays a role in the anxiety and microglial function rescue observed with soil intervention. After three generations of antibiotic exposure in mice, the abundance of functional genes involved in bacterial secretion systems, flagellar assembly, and two-component systems in adult G3 mice decreased significantly, and restored by soil intervention to a large extent. Most of these functional genes are present in pathogenic bacteria, which can aid in their ability to infect humans[43]. For instance, Helicobacter pylori contains a type IV secretion system and is known to cause gastritis and gastric ulcers. However, recent research by Blaser et al. [44]suggests that the absence of Helicobacter pylori can lead to acid reflux and other illnesses. The role of these conditional pathogenic bacteria and their secretion systems in human immune stimulation is not well understood.

The mechanisms underlying soil rescue may be related to the close contact between humans and soil microbes during the co-evolution of humans and their gut microbes. Humans used to breathe air containing soil aerosols on a daily basis, eat unwashed plant roots stuck with soil, or even consciously consume some products containing soil [45-47]. However, with the development of social civilization, human life has become increasingly isolated from soil. Therefore, soil supplement may now support the growth of key intestinal microorganisms and achieve the goal of restoring intestinal microecology.

In conclusion, this study simulated an important process of the gradual disappearance of modern human gut microbes and demonstrated that soil can effectively reverse this process as a prebiotic. This suggests that the occurrence of multiple modern human chronic diseases is associated with antibiotic use and soil rescue is an effective means of addressing these issues. The study provides a theoretical and experimental basis for the research and treatment of multiple modern chronic diseases related to intestinal microorganisms.

## Materials and methods

### Animals

The mice in the experiment shared a similar living environment and genetic background. All of them were provided with sterile drinking water. The only difference was that the mice receiving soil intervention were given soil as a supplement, while the others were fed the same sterile diet. The bedding materials for all mice were sterile, and their cages were kept at a temperature of 24 ± 2 °C and a humidity of 40% ± 5%, with a 12-hour light/dark cycle. The bedding was changed once a week, and all cages were covered with breathable mesh to ensure that microorganisms could spread through the air and between different cages.

#### Experimental model

Four-week-old SPF C57/BL6 male and female mice were treated with vancomycin for one week, followed by streptomycin for one week, and a metal mesh was used to achieve fecal-oral separation (Fig. S22). After 4-8 weeks of antibiotic withdrawal, the mice were bred, and the dams were changed to cages with normal bedding at approximately 15 days after pregnancy until the offspring were separated. The resulting mice were named G1’ mice. The same antibiotic and breeding treatment was performed on G1’ mice to obtain G2’ mice. G3’ mice were obtained in the same way (Fig. 1A).

#### Experiments on adult mice

The adult mice experiment involved G0’ mice, which were SPF mice. At 5 weeks of age in the SPF (G0’), G1’, G2’, and G3’ groups, we transferred the four groups to a general animal house with abundant environmental microorganisms. After they had been in the new environment for 2 weeks, fecal samples were collected and named G0, G1, G2, and G3 groups, respectively. The other half of the G0’, G1’, G2’, and G3’ groups were moved to the general animal house and fed a diet containing 5% sterilized soil. After two weeks of soil rescue, fecal samples were collected and named G0R, G1R, G2R, and G3R groups, respectively (Fig. 1A and Table S9). The collected feces were frozen at -80 [for further analysis of the intestinal microbiota. Brain and colon samples were also collected (refer to supplementary materials).

#### Experiments on infant mice

On the second day after birth, we transferred the SPF and G3’ groups and their dams from SPF animal room to the general animal room, which was rich in environmental microorganisms, to obtain the PG0 and PG3 groups. The infant mice from both the SPF and G3’ groups, along with their dams, were transferred to the general animal room, and they were fed daily with sterilized soil to obtain the PG0R and PG3R groups (Fig. 4A and Table S9). Specifically, the infant mice were fed with 10 μL of soil suspension (0.5 g/mL in water) daily for the first three days, followed by 20 μL of soil suspension daily for the next seven days. After 10 days of being transferred to the general animal house, we collected the rectum and contents of the mice, stored them in 1 mL of normal saline, and froze them at -80[for further analysis of the intestinal microbiota. We also collected the brain and colon for analysis (refer to the supplementary materials)

### Soil rescue for adult and infant mice

Soil was collected from the farm floor at a depth between 0.5 cm and 10 cm [24]. The collected soil was then sterilized using a 121°C autoclave for 30 minutes, repeating the process three times with 24-hour intervals in between. The sterilized soil was stored at -20°C until further use. For adult mice, the sterilized soil was crushed and mixed with their diet at a ratio of 5/95. For infant mice, the sterilized soil was mixed with sterile water at a mass ratio of 1/2 to form a paste for intervention.

## Data availability

Metagenomic sequence data for each mouse are available from the Sequence Read Archive (accession number: PRJNA947350).

## Acknowledgements

This work was supported by The Natural Science Foundation of China (grant no. 31770540) and The Key Research Program of Jiangsu (grants no. BE2018663).

## Author Contributions

Conceived and designed the experiments: NL, DZ.

Animal experiments: NL, HZ, YD, ML, DZ.

Related experiments and data analysis of gut microbiota and behavior: NL, HZ, ZB, YD, ML, WZ, ZF, XS, PX, DZ.

Experimental design related to microglia and intestinal barrier: YG.

Related experiments and data analysis of microglia and intestinal barrier: XX, YG, JS.

Writing of microglia and intestinal barrier: XX, YG. Writing of other sections: NL, DZ.

