## Supplementary figure S1-S22 for "Sterile Soil as Prebiotics Mitigate Intergenerational Loss of Gut Microbial Diversity and Anxiety-Like Behavior Induced by Antibiotics in mice"

**This file includes:**

Figures. S1 to S22

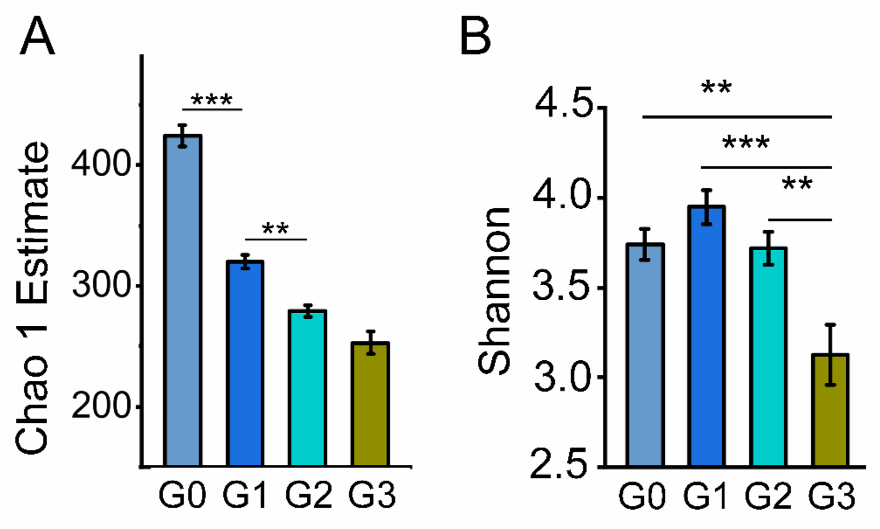


Figure S1 Effect of antibiotic treatment on gut microbial diversity in three generations of mice. A and B. Chao 1 estimate and Shannon diversity index of faecal micrbiota from each generation of mice. (n = 9 or 10; Error bars are s.e.m. and *P* values are based on ANOVA with bonferroni post-hoc test. *$P$ < 0.05, **$P$ < 0.01, ***$P$ < 0.001).

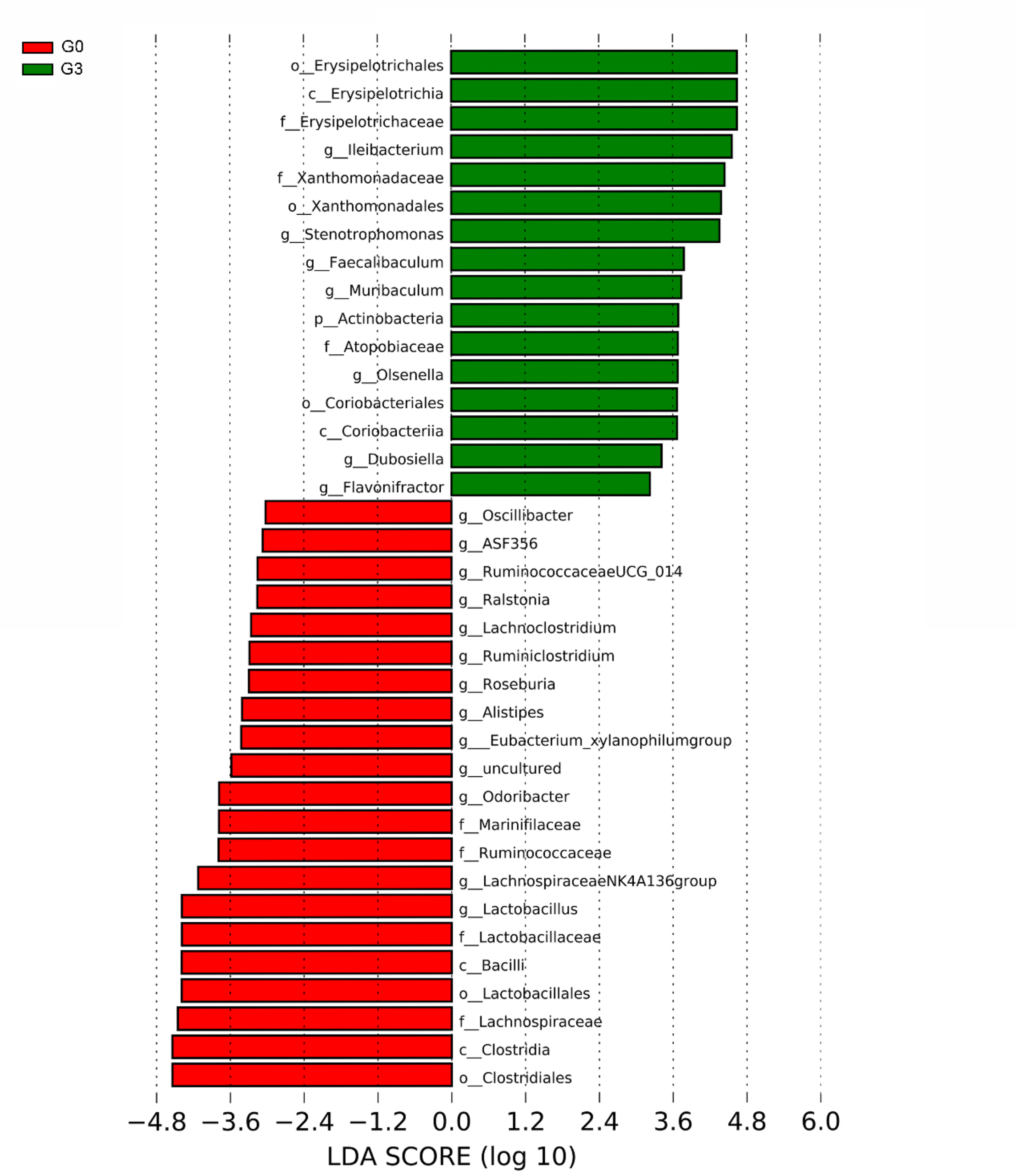


Figure S2. LEfSe Analysis on the Taxa of G0 and G3 Groups (LDA score>3).

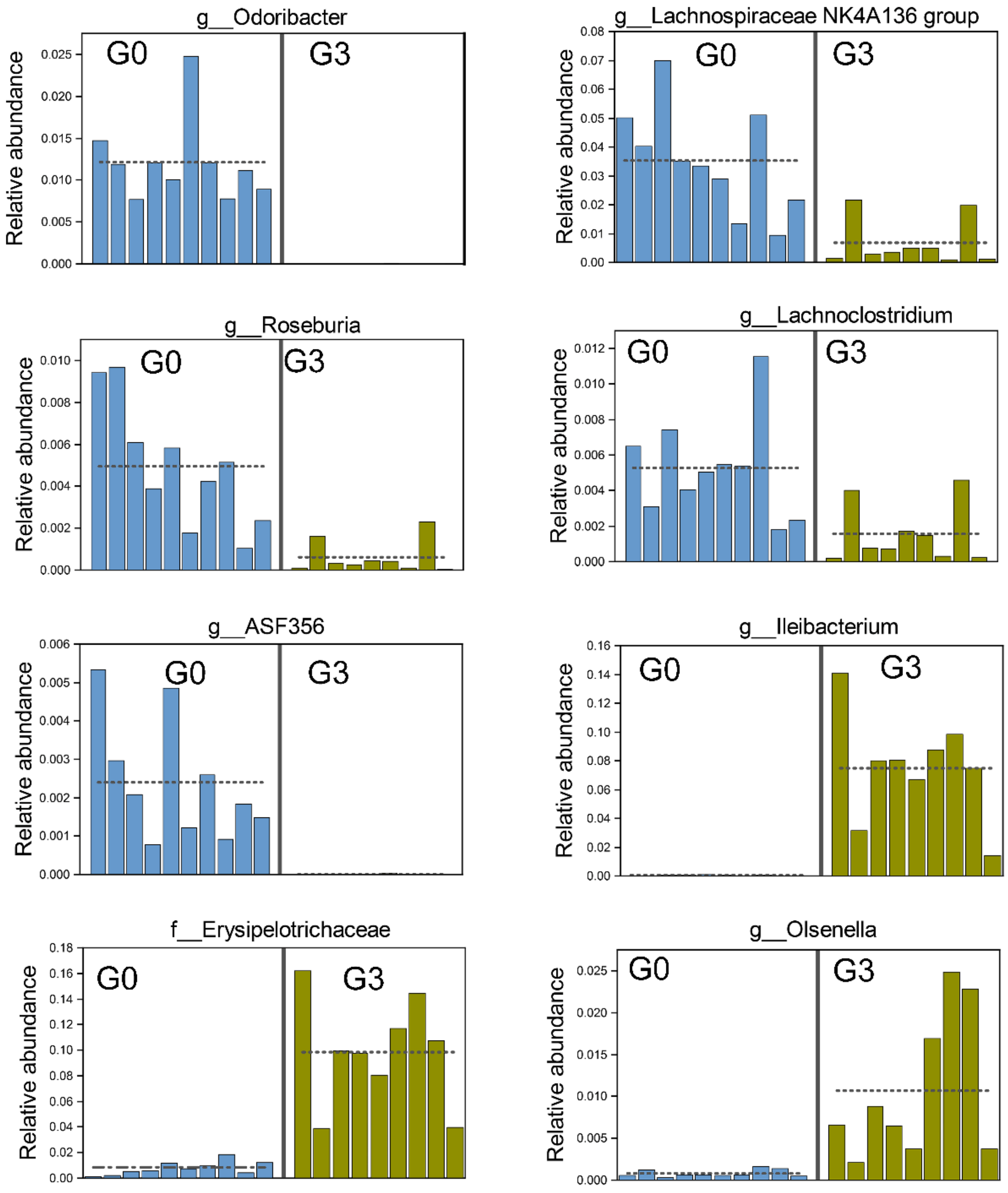


Figure S3. Histograms of some taxa with LDA>3 in lefse analysis. One column in the figure represents a sample.

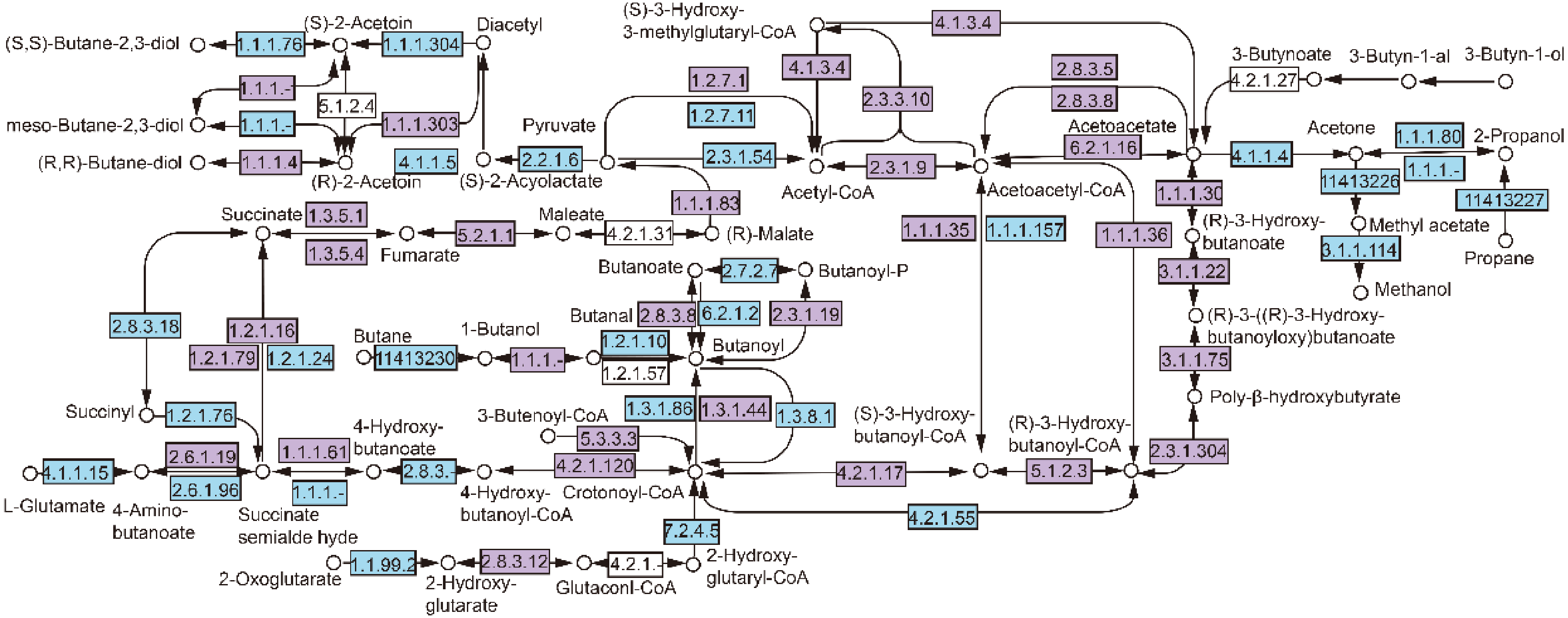


Figure S4. Functional genes associated with butyric acid metabolism. The purple box represents G0 highly expressed genes (P value of Mann-Whitney U test is less than 0.05 and the fold change of the two groups is greater than 2), and the blue box represents genes without difference (P value of Mann-Whitney U test is greater than or equal to 0.05 or the fold change of two groups is less than 2) or undetected.

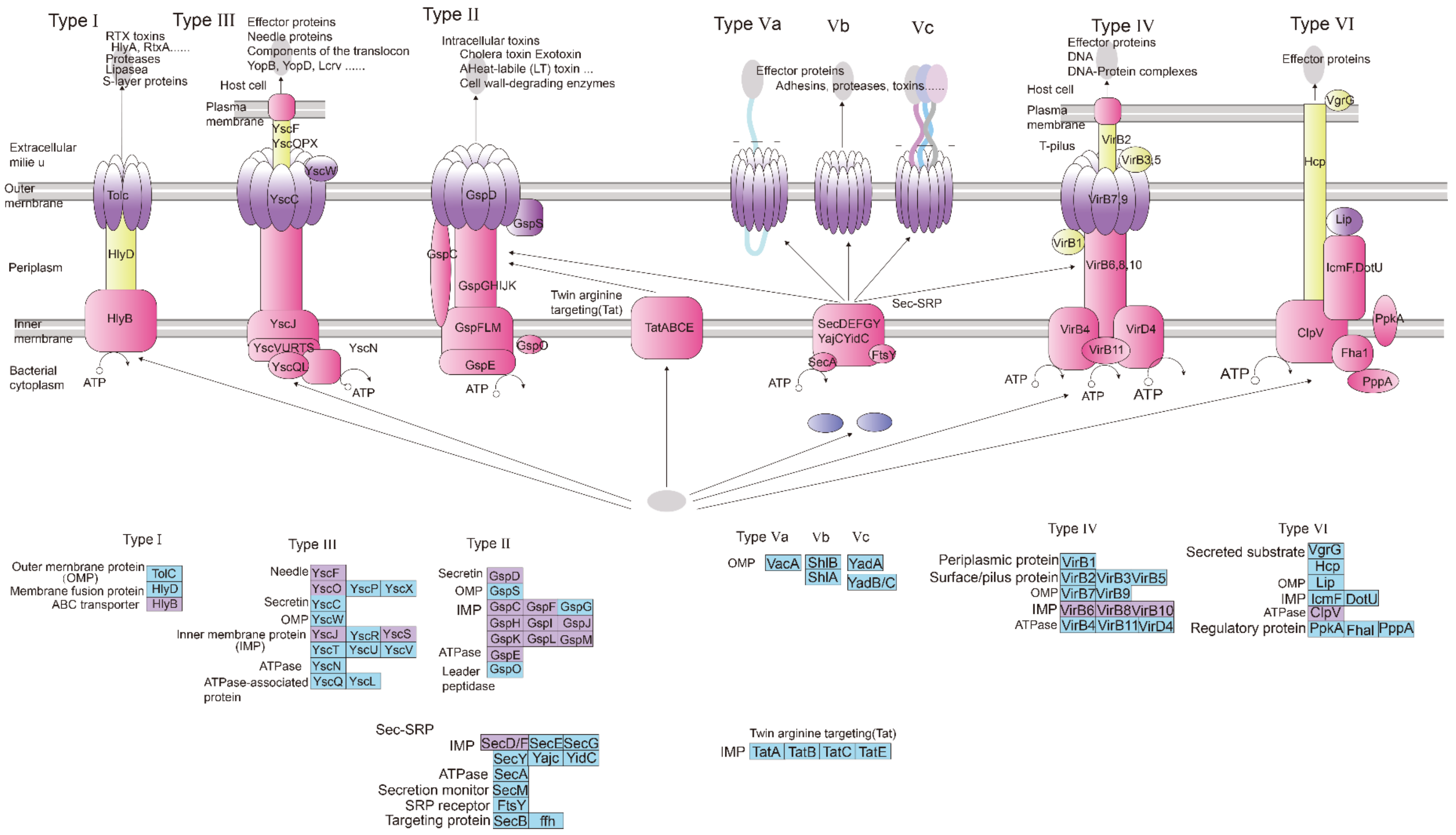


Figure S5. Functional genes associated with bacterial secretion system. The purple box represents G0 highly expressed genes (P value of Mann-Whitney U test is less than 0.05 and the fold change of the two groups is greater than 2), and the blue box represents genes without difference (P value of Mann-Whitney U test is greater than or equal to 0.05 or the fold change of two groups is less than 2) or undetected.

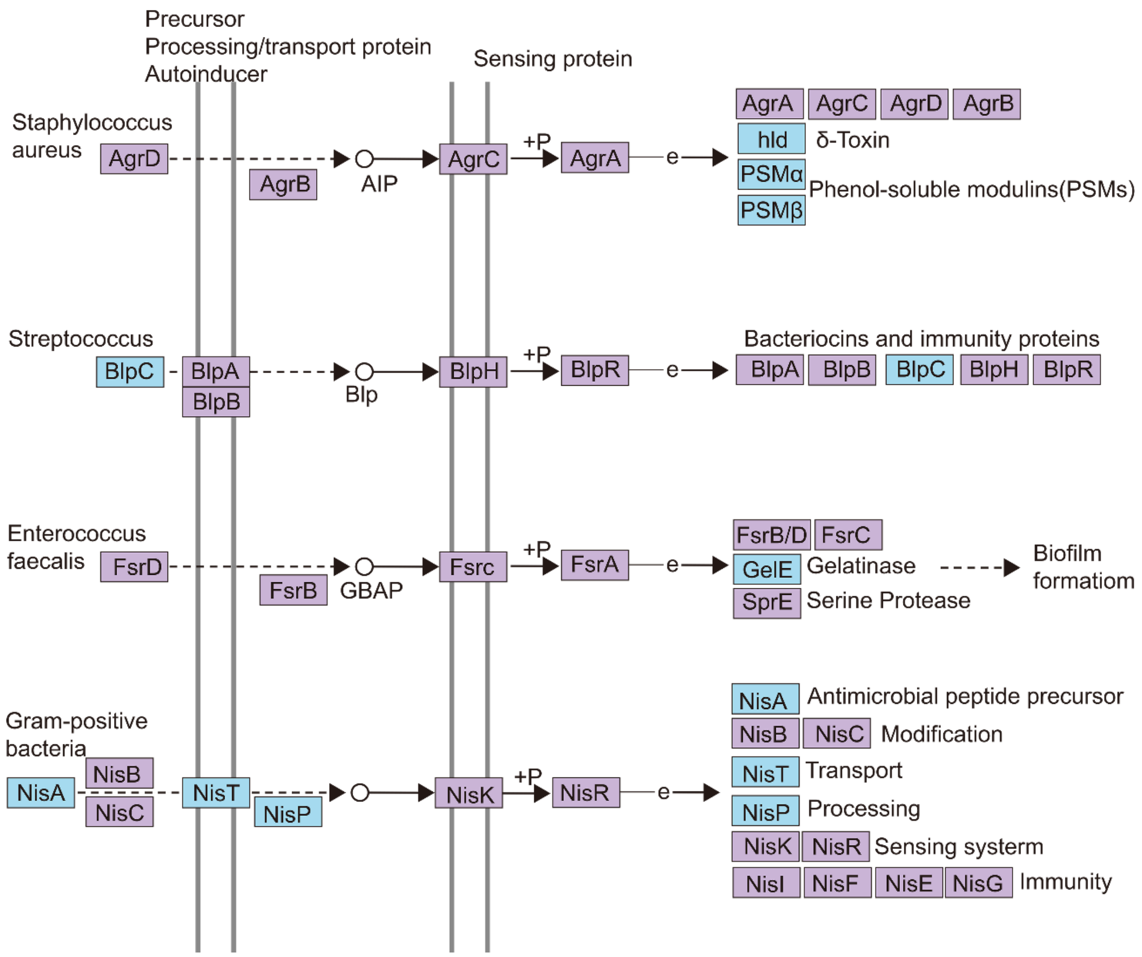


Figure S6. Functional genes associated with Two-component system. The purple box represents G0 highly expressed genes (P value of Mann-Whitney U test is less than 0.05 and the fold change of the two groups is greater than 2), and the blue box represents genes without difference (P value of Mann-Whitney U test is greater than or equal to 0.05 or the fold change of two groups is less than 2) or undetected.

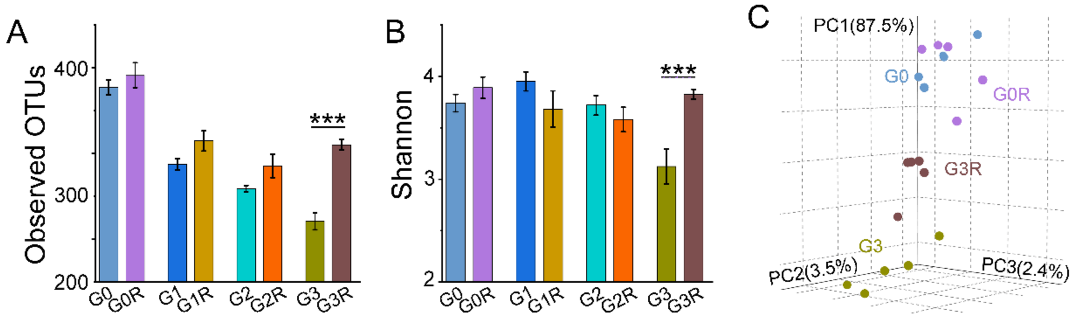


Figure S7. The repair effect of soil rescue on intestinal microecological dysbiosis in the offspring of parental mice using antibiotics. A and B. Number of OTUs at a 97% threshold and Shannon diversity index of faecal microbiota from each generation of mice. (n = 9 or 10; Error bars are s.e.m. and *P* values are based on ANOVA with bonferroni post-hoc test. $*P < 0.05$, $**P < 0.01$, $***P < 0.001$). C. Principle coordinates analysis (PCoA) of bray distance of shotgun genes.

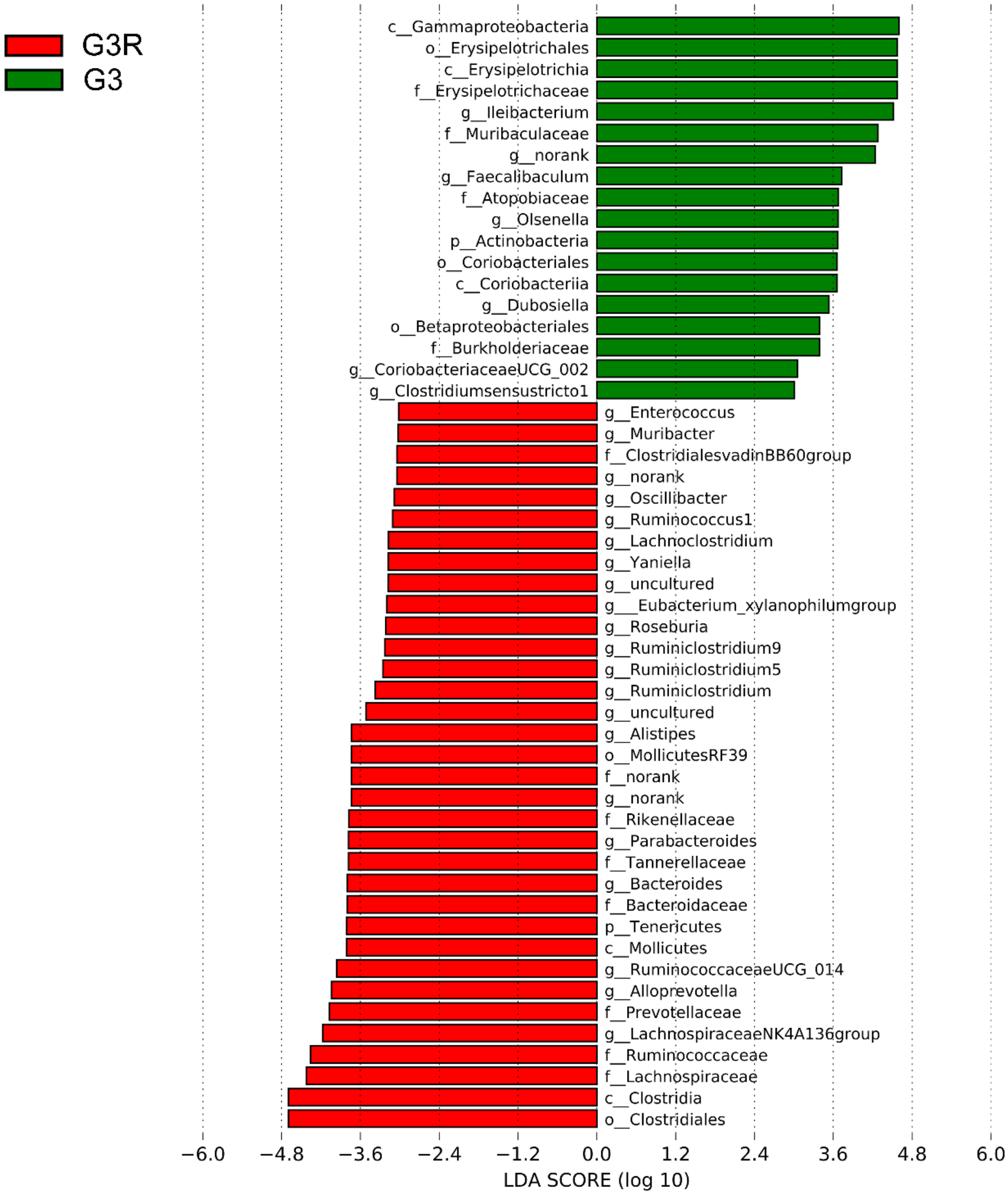


Figure.S8. LEfSe Analysis on the Taxa of G3 and G3R groups (LDA score>3).

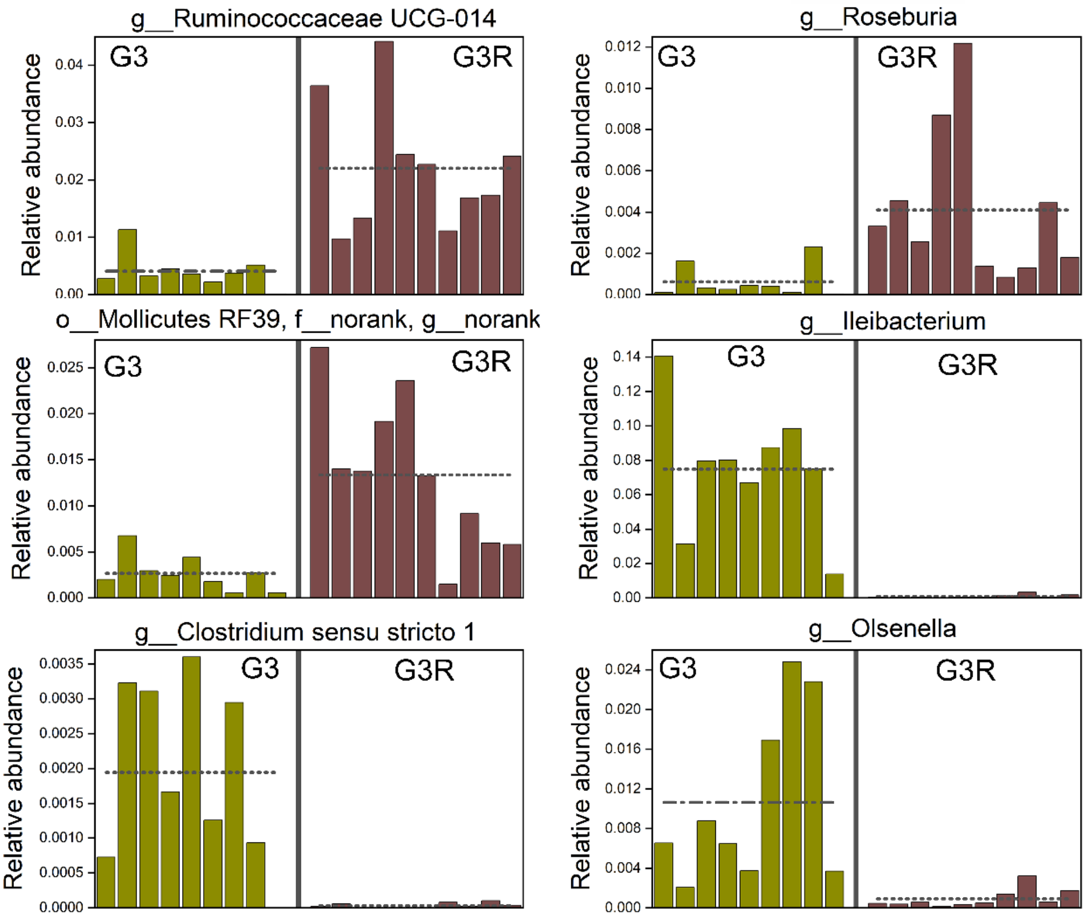


Figure. S9. Histograms of some taxa with LDA>3 in lefse analysis. One column in the figure represents a sample.

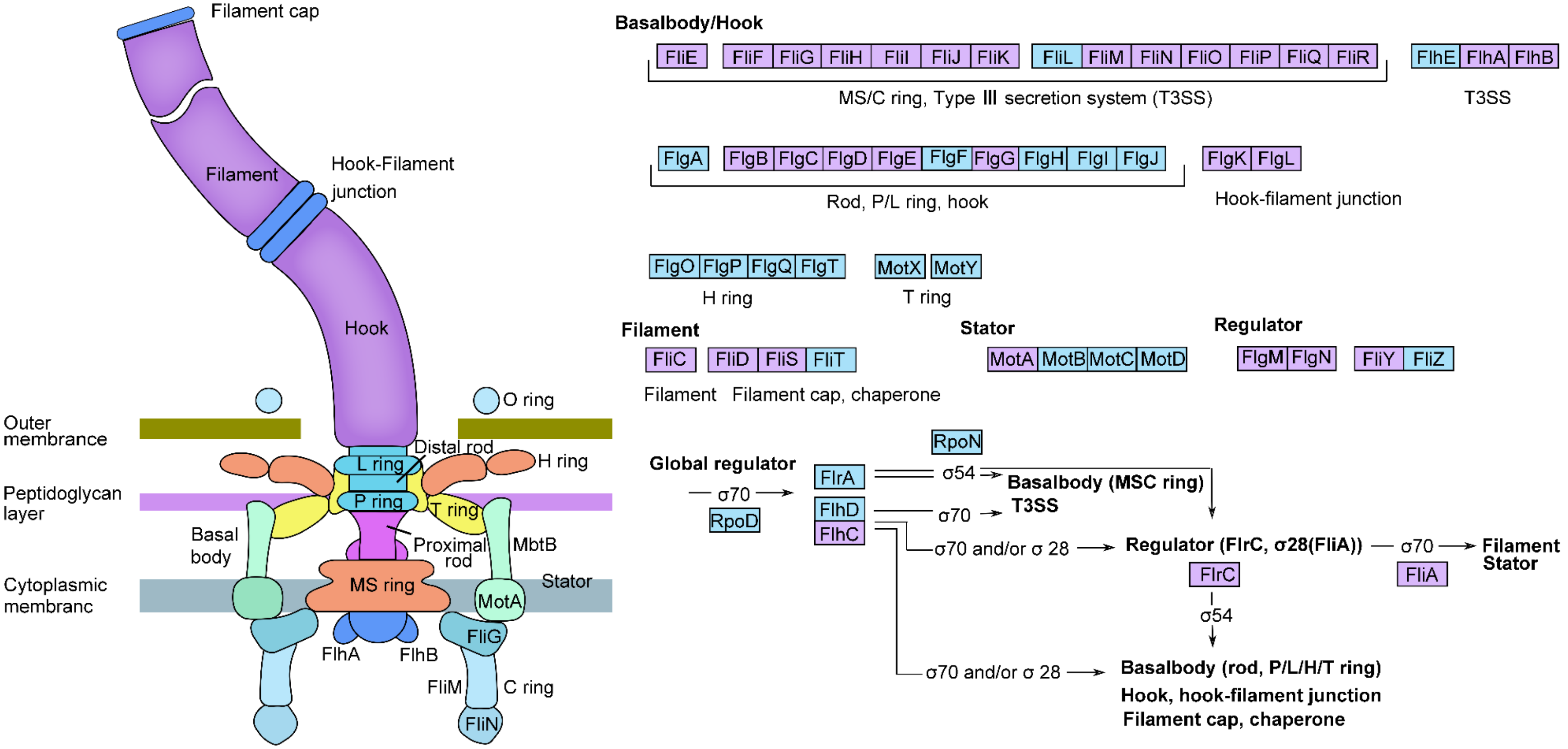


Figure S10. Functional genes associated with flagellar assembly. The purple box represents G3R highly expressed genes (P value of Mann-Whitney U test is less than 0.05 and the fold change of the two groups is greater than 2), and the blue box represents genes without difference (P value of Mann-Whitney U test is greater than or equal to 0.05 or the fold change of two groups is less than 2) or undetected.

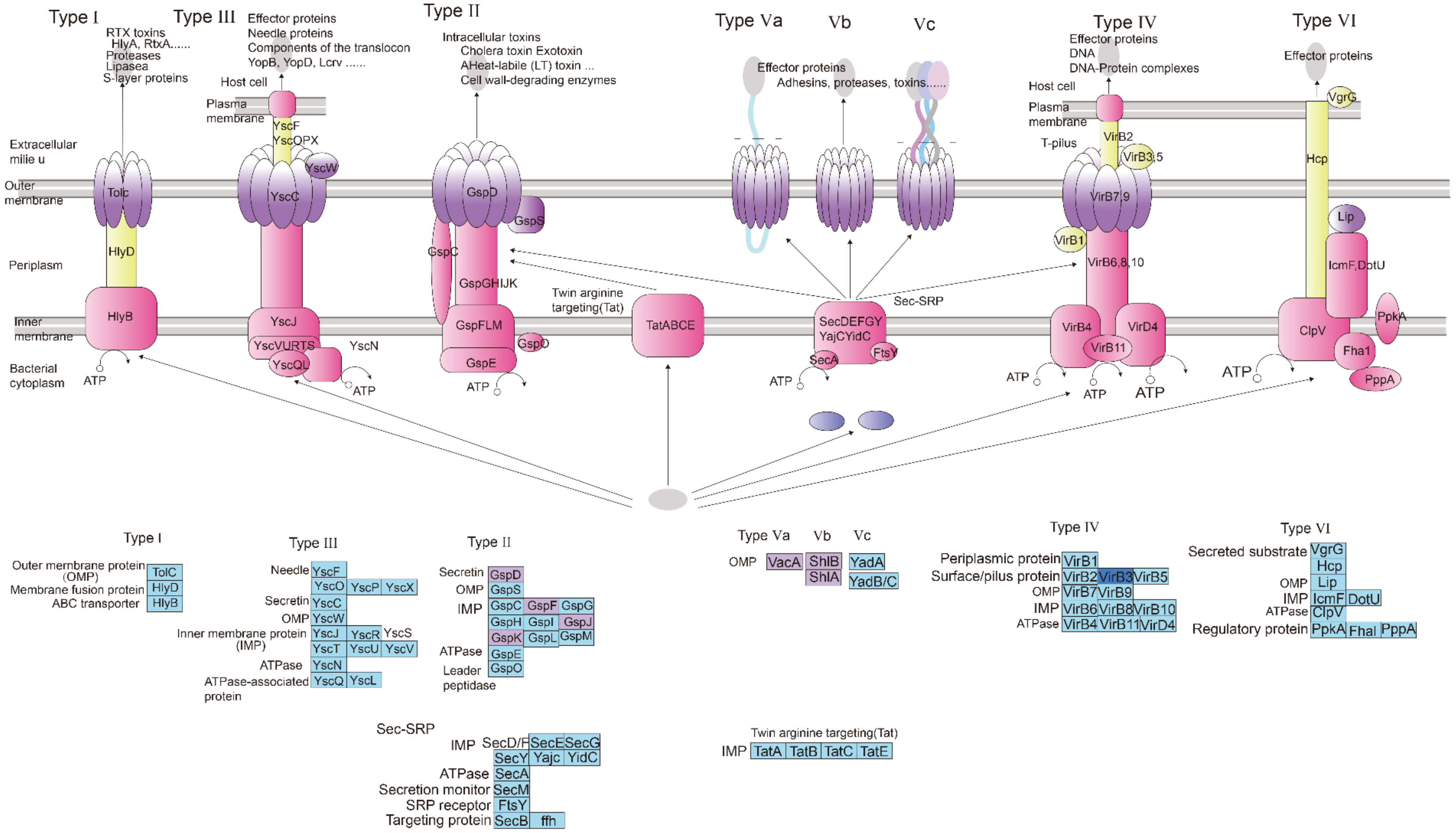


Figure S11. Functional genes associated with bacterial secretion system. The purple box represents G3R highly expressed genes (P value of Mann-Whitney U test is less than 0.05 and the fold change of the two groups is greater than 2), the blue box represents genes without difference (P value of Mann-Whitney U test is greater than or equal to 0.05 or the fold change of two groups is less than 2) or undetected, and dark blue represents high expression in the G3R group.

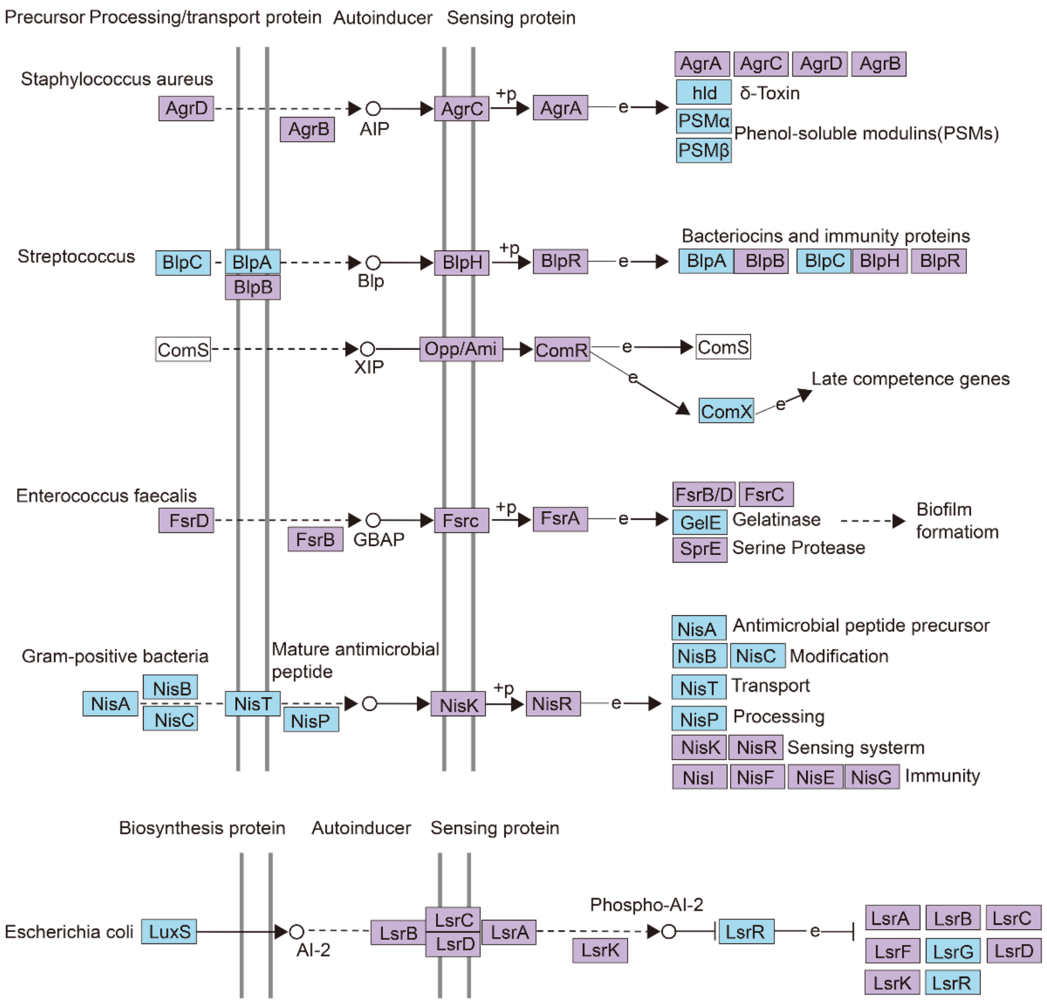


Figure S12. Functional genes associated with Two-component system. The purple box represents G3R highly expressed genes (P value of Mann-Whitney U test is less than 0.05 and the fold change of the two groups is greater than 2), and the blue box represents genes without difference (P value of Mann-Whitney U test is greater than or equal to 0.05 or the fold change of two groups is less than 2) or undetected.

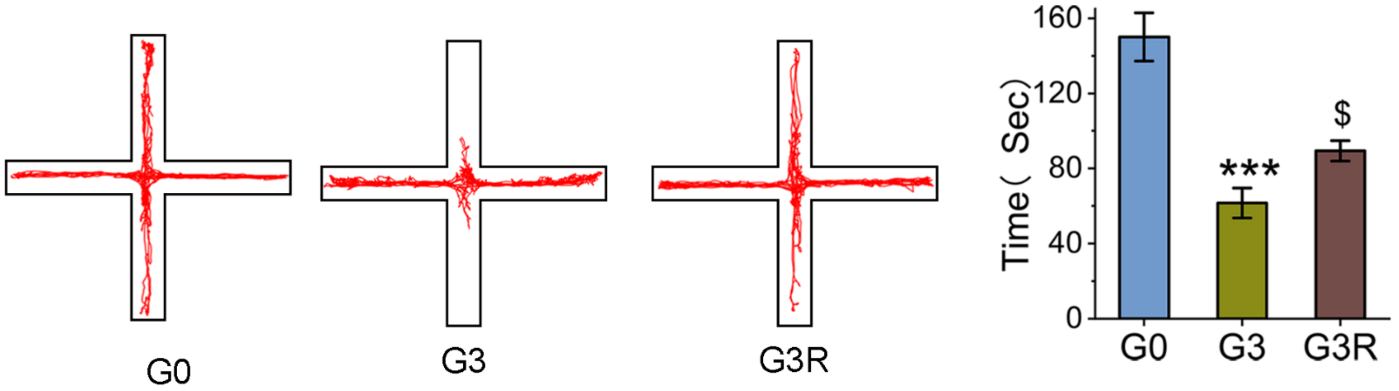


Figure S13. Elevated plus-maze test. (n=5, Error bars were s.e.m. and *P* values were based on 2-tailed t test. ***$P < 0.001$ vs G0, $^{\$}P < 0.05$ vs G3).

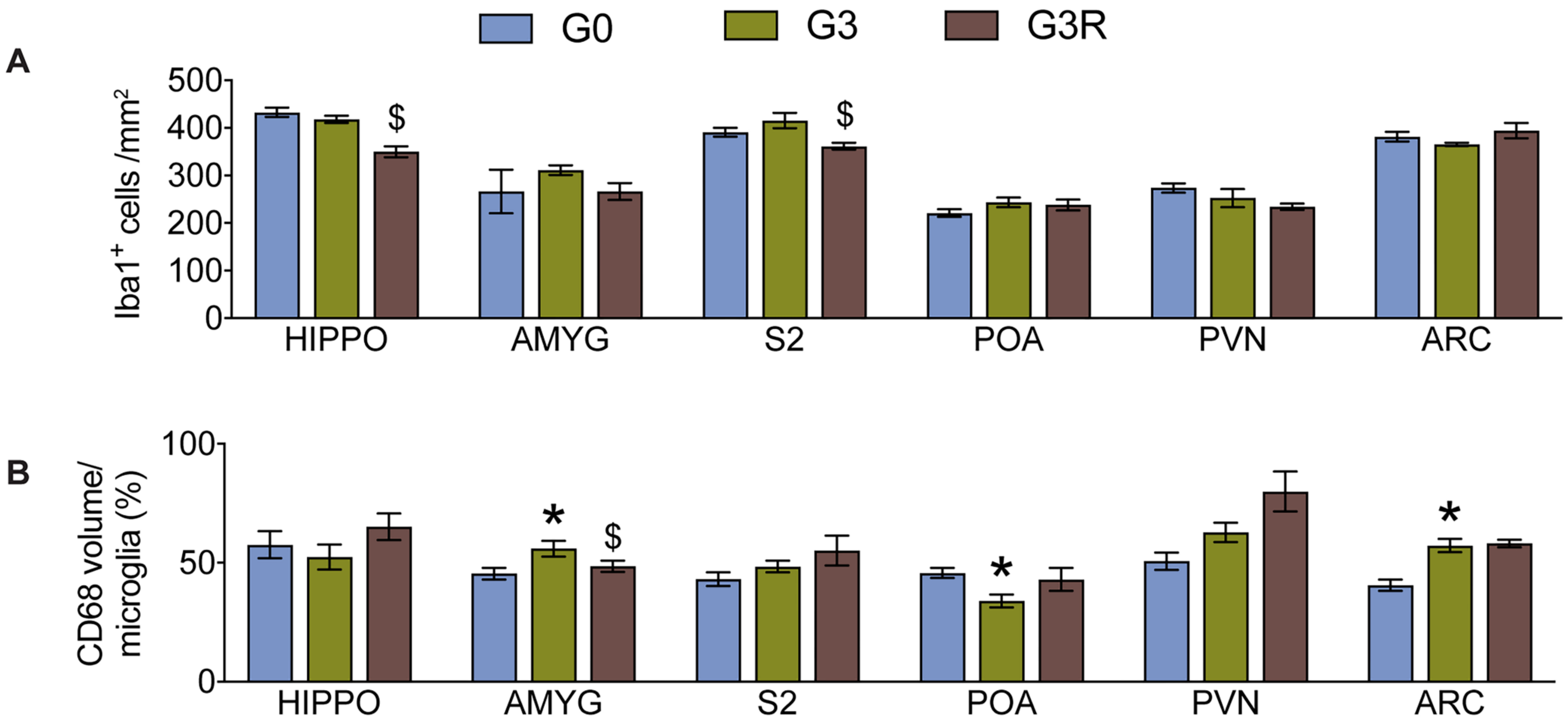


Figure S14. A. Quantification of Iba1-ir postivie cell number in each brain region. B. Quantification of CD68-ir positive volume in individual microglia in in each brain region. N=4-5 animals per group. *P<0.05 vs G0; $p<0.05 vs G3. All the mice in this figure are males.

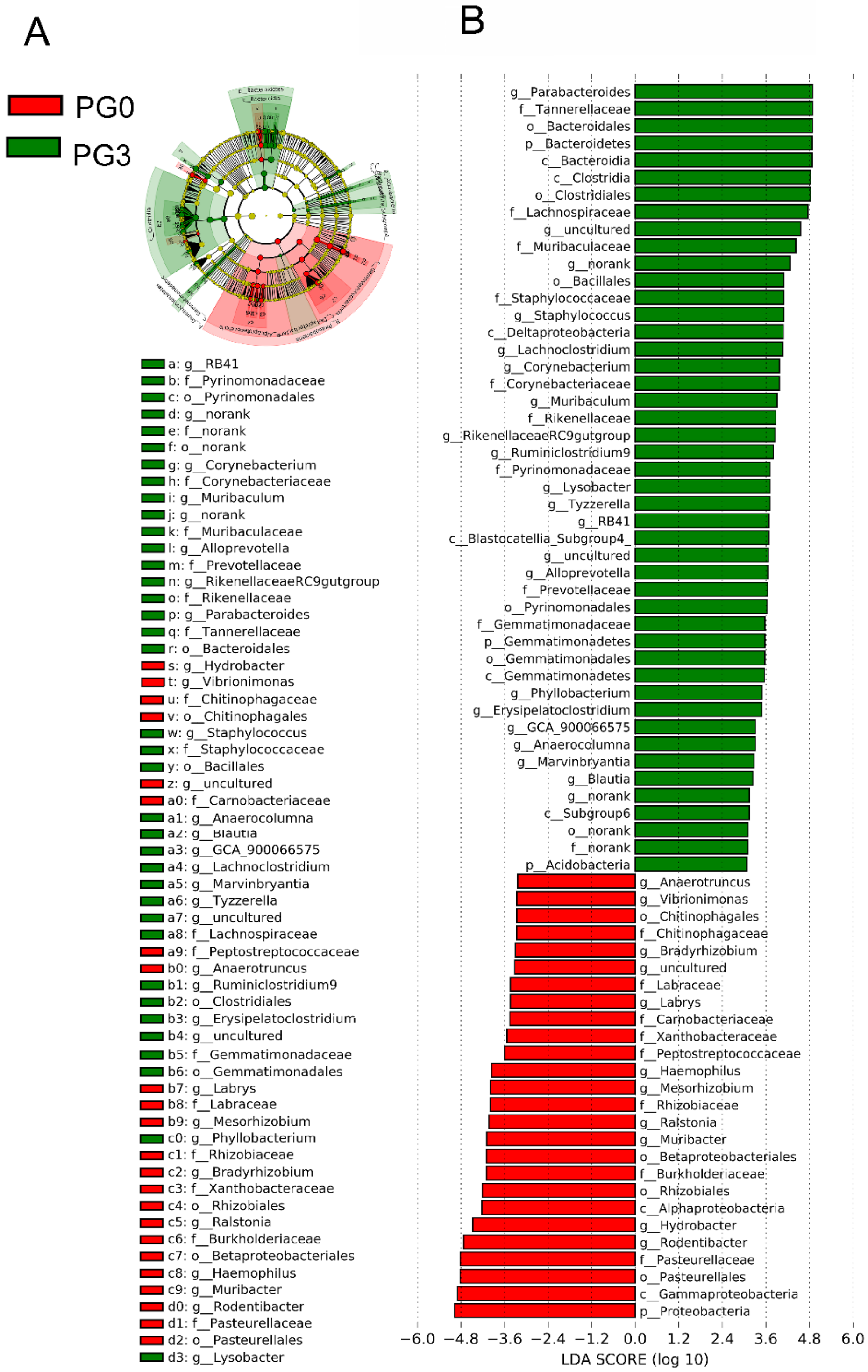


Figure S15. LEfSe Analysis on the Taxa of PG3 and PG3R Groups (LDA score>3).

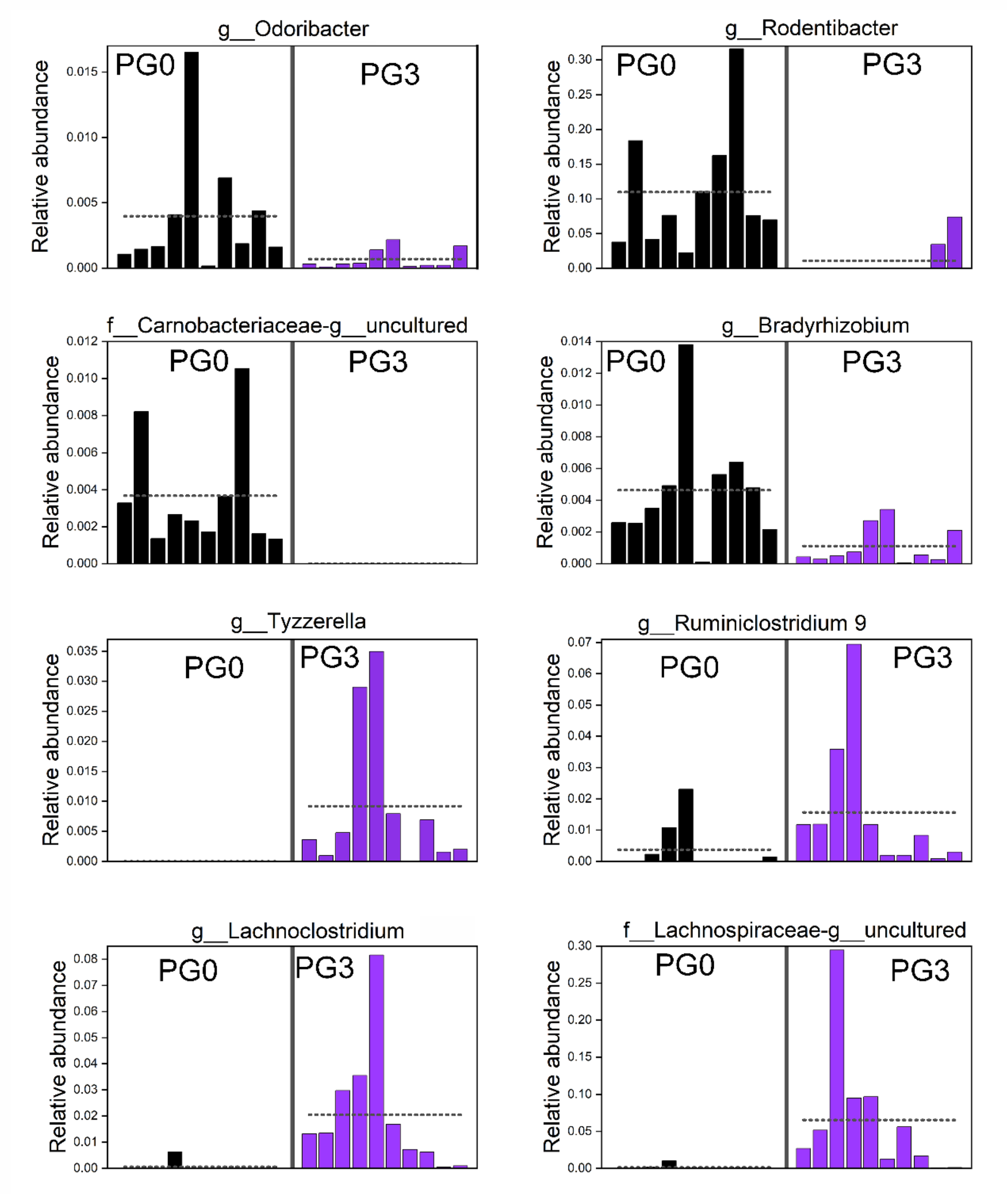


Figure S16. Histograms of some taxa with LDA>3 in lefse analysis. One column in the figure represents a sample

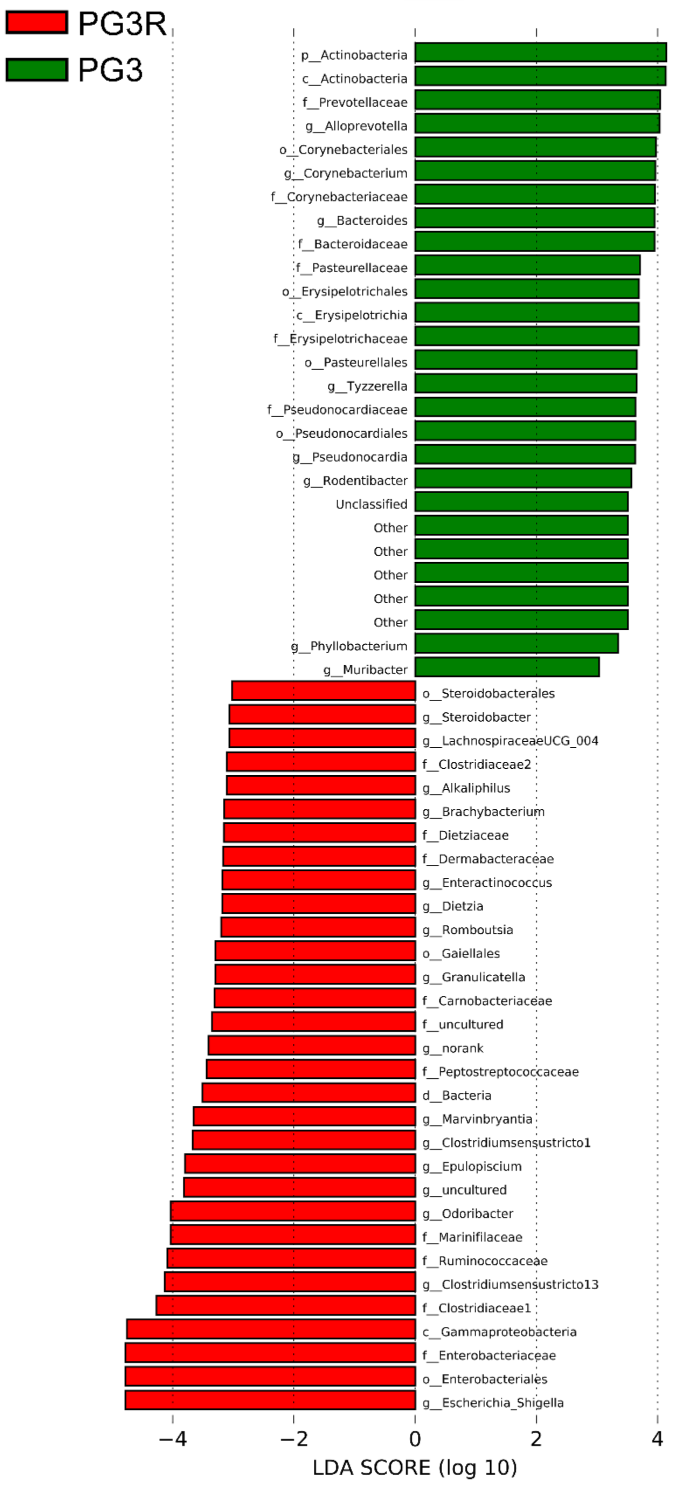


Figure S17. LEfSe Analysis on the Taxa of PG3 and PG3R Groups (LDA score>3).

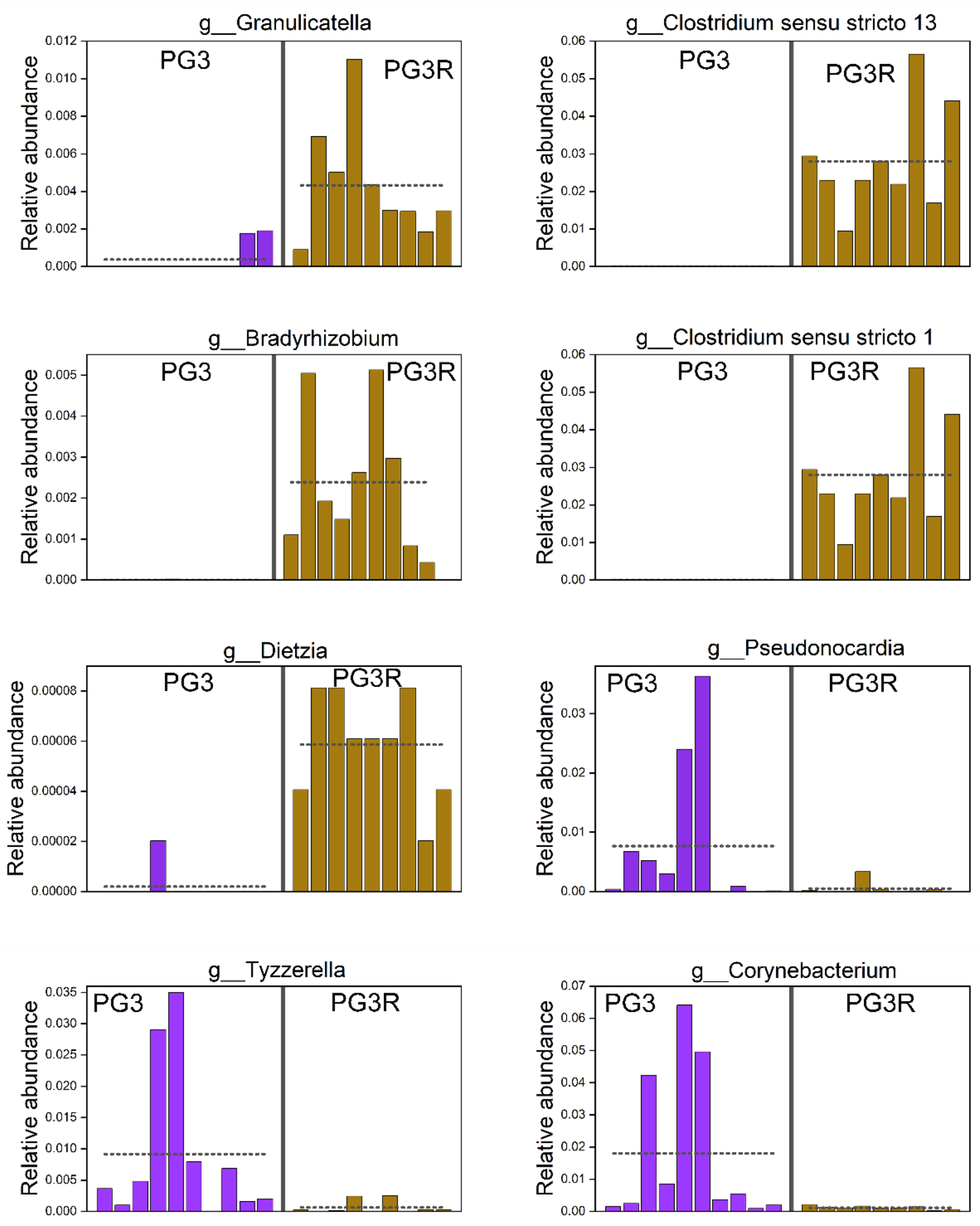


Figure S18. Histograms of some taxa with LDA>3 in lefse analysis. One column in the figure represents a sample.

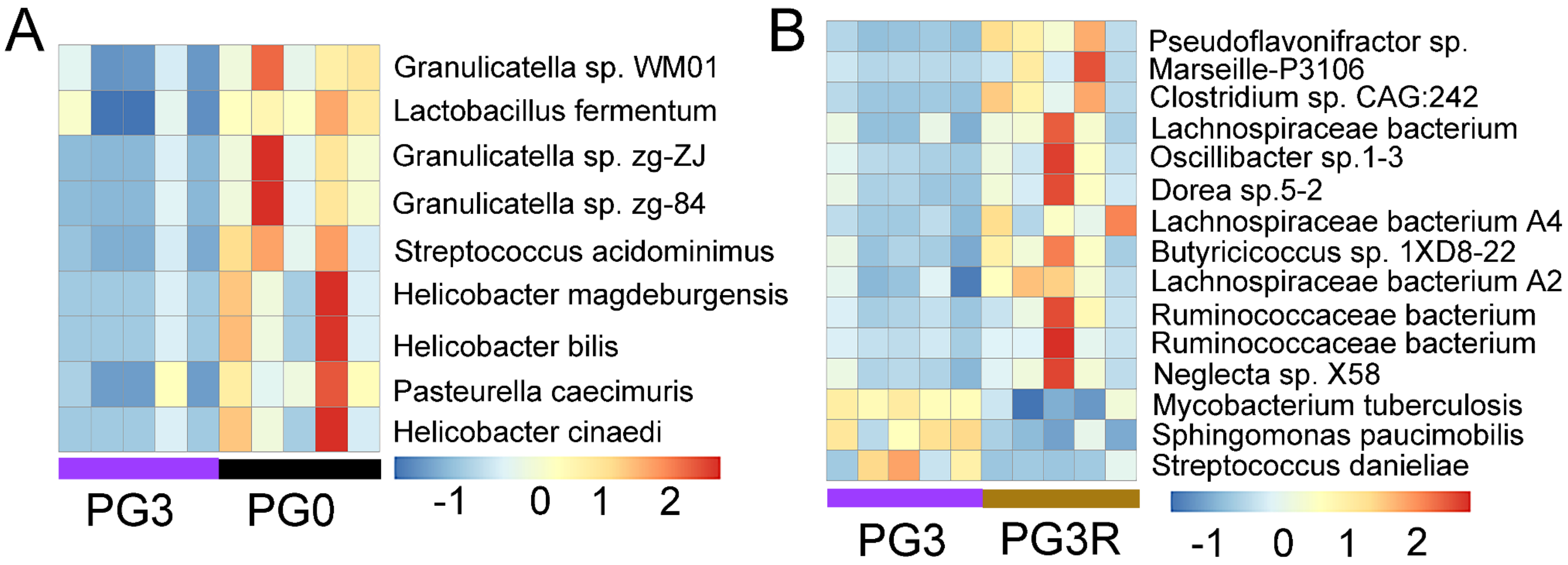


Figure S19. A. Significantly different species among top 500 species between PG3 and PG0 groups. B. Significantly different species among top 500 species between PG3 and PG3R groups. The species showed in figure A and B with P value of Mann-Whitney U test less than 0.05. The color showed the Z score of relative abundance. Data was based on shotgun sequences.

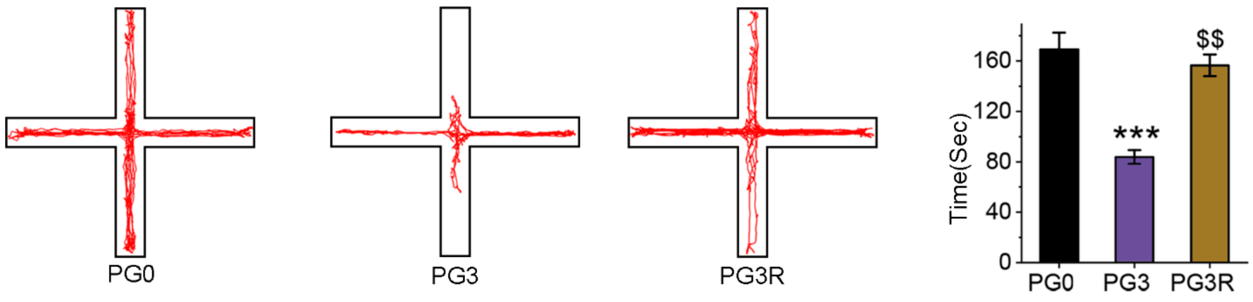


Figure S20. Elevated plus-maze test. (n=5, Error bars were s.e.m. and P values were based on 2-tailed t test. ***P < 0.001 vs PG0, $P < 0.05 vs PG3).

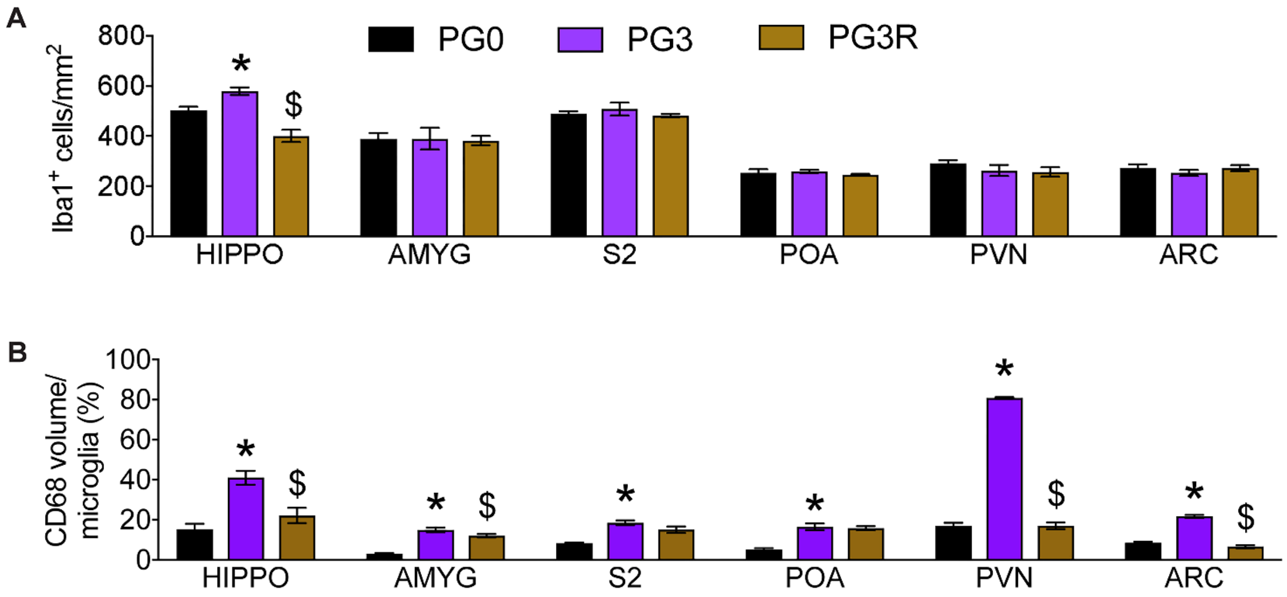


Figure S21. A. Quantification of Iba1-ir postivie cell number in each brain region. B. Quantification of CD68-ir positive volume in individual microglia in in each brain region. N=4-5 animals per group. *P<0.05 vs G0; $p<0.05 vs G3. All the mice in this figure are males.

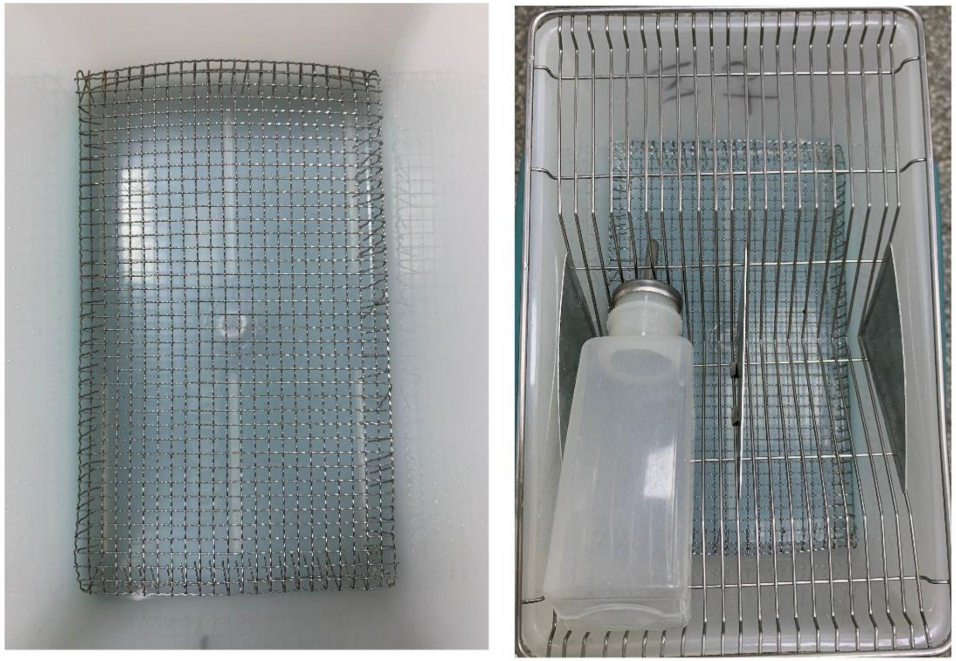

Figure S22. A cage with metal mesh for leaking feces.
