## Supplementary materials and methods for "Sterile Soil as Prebiotics Mitigate Intergenerational Loss of Gut Microbial Diversity and Anxiety-Like Behavior Induced by Antibiotics in mice"

**Care and use of experimental mice**

Animal experiments were conducted in strict accordance with the guidelines provided by the Animal Research Ethics Committee of Southeast University. All animal experiments were approved by the Animal Care Research Advisory Committee of Southeast University and the National Institute of Biological Sciences (approval number: 2017063009). We made every effort to minimize and alleviate the suffering that the animals might experience. We monitored the health of the mice every other day and weighed them once a week. The health status of the mice was evaluated by observing changes in their body weight and fecal shape and appearance. In the event of severe illness, including weight loss of 15% to 20%, diarrhea, hair loss, arched or curled posture, or lethargy lasting more than a week, the animals were euthanized to minimize pain and suffering. None of the mice displayed any symptoms during the experiment.

**Behavioral experiments**

To acclimate the mice to the testing environment, we moved the experimental mice to the testing room 24 hours before the tests. Before testing each animal, we cleaned the testing equipment with 75% ethanol to prevent any potential odors left by other mice from interfering with the tests. All behavioral tests were conducted during the photoperiod, and the behavior of the mice was recorded and blindly analyzed using Noldus Ethovision XT software.

After two weeks of being housed in the general animal room, we stopped the soil rescue in the G3R group and tested the behavior of the mice in the G0, G3, and G3R groups. Similarly, ten days after moving the infant mice to the general animal house, we stopped soil intake in the PG3R group. The PG0, PG3, and PG3R mice were kept in the general animal room until they reached six weeks of age, after which their behavior was tested.

**Open-Field (OFT) Test:** We placed each mouse in a plastic chamber (50 × 50 × 50 cm) with an opening at the top and opaque on all other sides. The mouse was then allowed to explore the chamber freely for five minutes. We recorded the time and frequency of the mice entering the central area of the chamber, which accounted for half of the total area.

**Elevated plus maze:** The elevated plus maze comprises of two open arms and two closed arms of equal size. Each arm is 35 × 5 cm in size, with walls that are 15 cm in height. The maze is made of white plastic panels and is elevated 60 cm from the floor. To conduct the experiment, we placed the mice in the center of the maze, facing one of the open arms, and allowed them to explore freely for five minutes. We recorded the time when the mice entered the open arm.

**16S rDNA sequencing**

To extract gut microbial DNA, we used the E.Z.N.A.® Stool DNA Kit (Omega Bio-tek, Norcross, GA, USA) and amplified the V4 region of the 16S rDNA using primers 515F and 907R [1]. The amplified products were purified using the AxyPrep DNA Gel Extraction Kit (Axygen Biosciences, Union City, CA, USA) and quantified using QuantiFluor™-ST (Promega, USA). We then pairedend sequenced (2 × 250) the PCR product library using the Illumina MiSeq platform. To ensure data quality, we removed low-quality sequences using Trimomatic (version 0.32) [2] and chimeric sequences using UCHIME (version 4.2.40) [3]. We used Usearch (version 7.1; http://drive5.com/uparse/) to define the OTUs with a threshold of 97% similarity, and the Ribosomal Database Project (RDP) Classifier (version 2.2; http://sourceforge.net/projects/rdpclassifier/) to determine the taxonomic identities of the phylotypes with a confidence threshold of 70%. The reads of each sample are presented in table S9. Shotgun metagenomic sequencing was also performed.Shotgun metagenomic sequencing

Metagenomic shotgun sequencing libraries were created and sequenced. Firstly, 1 μg of genomic DNA from each fecal sample was sheared into fragments approximately 450 bp in length using the Covaris S220 Focused-ultrasonicator (Woburn, MA, USA). These fragments were then used to construct sequencing libraries. The Illumina HiSeq X instrument was used to sequence the libraries in a pair-end 150-bp (PE150) mode. To remove adaptor contaminants and low-quality reads, Trimmomatic (version: 0.32; http://www.usadellab.org/cms/uploads/supplementary/Trimmomatic) was used[2]. The Burrows-Wheeler Aligner (BWA, version: 0.6) mem algorithm (parameters: -M -k 32 -t 16; http://bio-bwa.sourceforge.net/bwa.shtml) was used to map the human genome (version: hg19). Host genome reads were removed and clean reads were obtained. The taxonomy of the clean reads was then annotated using Kraken2 [4] with the customized Kraken database, and the abundance of the taxonomy was estimated using Bracken (version: 2.0.0; https://ccb.jhu.edu/software/bracken/).

For this analysis, the clean sequence reads were used to generate a set of contigs using MegaHit (version: 1.1.4) with "--min-contig-len 500" parameters [5] . Then, Prodigal (version: 2.6.3) [6] was used to predict open reading frames (ORFs) of the assembled contigs. The ORFs were clustered using CD-HIT (parameters: -n 9 -c 0.95 -G 0 -M 0 -d 0 -aS 0.9 -r 1) [7] to generate a set of unique genes. The longest sequence of each cluster was selected as the representative sequence of the unique-gene set. To determine the abundance of each gene in the total sample, the Salmon software [8] was used to obtain the read number of each gene. Finally, the unique-gene set was searched against the KEGG database using Blastx to identify proteins and retrieve their functional annotations.

**Data analysis and statistical tests**

This passage describes the data analysis methods used for 16S rRNA gene sequencing data in both adult and infant mice, as well as for shotgun metagenomic sequencing data. Alpha diversity and beta diversity analysis of the 16S rRNA gene data were performed using the QIIME software package version 1.9.1. The rarefaction level for adult mice samples was 49265 sequences and for infant mice samples was 49124 sequences. Beta diversity analysis of the shotgun metagenomic sequencing data was performed using R version 4.0.5. Non-parametric statistical tests for OTU/species/KO were performed using R version 4.0.5, while other tests were performed using SPSS version 1.8.0.

**Immunohistology**

Mice were euthanized by isoflurane overdose and then perfused transcardially with saline. Brain and colon were fixed in 4% paraformaldehyde (PFA) for 48 h and dehydrated by 30% sucrose solution for 48h at 4℃. 30um coronal brain sections or 8um colon cross sections were pretreated with 10% H2O2 and 10% methanol, and then incubated with primary antibody overnight at 4°C, subsequently incubated with biotin-conjugated secondary antibody for 1 h at room temperature. For DAB staining, brain sections were incubated with avidin-biotin-horseradish peroxidase complex (1:800, vector, PK-6100), followed by 1% diaminobenzidine with 0.01% hydrogen peroxide for visualization. For immunofluorescent staining, brain sections were incubated with Alexa 488 or 555 fluorescent conjugated secondary antibody accordingly. All stained sections were imaged with a fluorescence microscope (Nikon, Melville, NY, USA). The following primary antibodies were used: anti-IBA1 antibody (rabbit, 1:800, Wako, 019-19741); anti-CD68 antibody (rat, 1:500, Abcam, 53444);

**Western blotting**

Colon tissues were rinsed in PBS to remove lumenal content and then homogenized in lysis buffer (Beyotime, Los Angeles, CA, USA) supplemented with cocktail protease inhibitor (Thermo Fisher Scientific, Waltham, MA, USA). 30 ug proteins were separated on 10% sodium dodecyl sulfate-polyacrylamide electrophoresis gel and transferred onto nitrocellulose membranes (Millipore). The membranes were blocked with 5% milk (Phygene, FuZhou, CN) for 1h at RT, and then incubated with the following primary antibodies overnight at 4°C: anti-β-ACTIN antibody (mouse, 1:500, ZSGB-Bio, TA-09)， anti-CLAUDIN-1 (mouse, 1:100; Santa Cruz, sc-166338), and anti-OCCLUDIN antibody (mouse, 1:100; Santa Cruz, sc-133256). The membranes were then incubated with HRP-conjugated secondary antibody for 1h at RT. ECL solution (Yeasen, 36208ES60) was used for visualization.
